# Dual pathway regulation of castration response and ferroptosis in the prostate epithelium

**DOI:** 10.1101/2025.08.06.668974

**Authors:** Weiping Li, Jing-Bo Zhou, Zejian Wang, Fereshteh Zandkarimi, Teresa A. Milner, Shouhong Xuan, John R. Christin, Caroline J. Laplaca, Andrew S. Greenberg, John P. Chute, Hanina Hibshoosh, Brent R. Stockwell, Michael M. Shen

## Abstract

Understanding how sex hormones maintain tissue integrity—and how their disruption can promote cancer—is a central question in biology. In the prostate, androgen signaling is crucial for development, homeostasis, and cancer, yet the molecular mechanisms underlying tissue regression after androgen-deprivation have remained unknown. Here we show that castration induces epithelial regression via ferroptosis in the normal mouse and human prostate, as well as in prostate tumors. Using *in vivo* analyses in genetically-engineered mice, supported by validation in human organotypic cultures, we demonstrate that androgen receptor (AR) signaling controls castration response through two distinct pathways: an intrinsic luminal epithelial pathway regulated by the prostate-specific transcription factor NKX3.1, and an extrinsic stromal signal mediated by the secreted factor pleiotrophin (PTN). Together, these AR signaling pathways coordinately regulate biosynthesis of monounsaturated fatty acid (MUFA) phospholipids and GPX4 expression to suppress prostate epithelial ferroptosis. Our findings reveal a sex hormone-regulated ferroptotic program that governs tissue homeostasis and suggest that ferroptosis induction could represent a new therapeutic strategy for prostate cancer.

## Introduction

Sex hormone signaling plays a vital role in the maintenance of secondary sexual tissues and its disruption can often promote cancer. In the case of the prostate, androgen signaling is essential for organogenesis and normal adult tissue homeostasis (*1, 2*). In the adult prostate, AR is widely expressed in both the stromal and epithelial compartments, and is believed to mediate stromal-epithelial interactions (*1-3*). Luminal cells are the most abundant epithelial cell type and represent a major target of castration-induced changes in the prostate epithelium (*3, 4*). Since both stromal and epithelial cells have active AR signaling, however, the specific contribution of individual compartments for castration response in luminal cells has remained unknown.

Over 80 years ago, Huggins and Hodges published their seminal finding that androgen-deprivation leads to regression of prostate tumors (*5*), which has provided the foundation for frontline prostate cancer therapy to the present day. Androgen-deprivation therapy (ADT) through chemical or surgical castration leads to regression of normal prostate tissue as well as hormone-sensitive prostate tumors, yet the molecular basis for this castration response has remained poorly understood. Even when prostate tumors recur following ADT in the form of castration-resistant prostate cancer (CRPC), these tumors usually remain dependent on AR activity, which has led to the development and clinical application of potent second-generation AR pathway inhibitors (ARPIs) such as enzalutamide (*6*). However, many tumors will ultimately develop resistance to ARPIs, leading to the emergence of aggressive AR-low or AR-negative subtypes through lineage plasticity (*6*). Thus, the development of new therapeutic approaches to improve the response to ADT as well as to overcome castration-resistance represents a critical unmet need.

Early work on castration response that was largely performed in the 1980s described decreases in tissue size as well as apoptosis in luminal epithelial cells following androgen-deprivation in the rodent and human prostate (*4, 7-9*). Further studies also indicated an essential role for stromal-epithelial interactions in castration response (*10*), but the key stromal secreted signal that mediates such interactions has remained unknown. In addition, the apoptotic index reported in these early studies is substantially lower than can account for the extensive cell death observed during regression, raising the possibility that a non-apoptotic form of regulated cell death could be involved, although such cell death mechanisms were not known at the time.

Ferroptosis was first described in 2012 as an iron-dependent form of regulated cell death due to lipid peroxidation (*11*). Subsequent studies have extensively elucidated mechanisms that govern ferroptosis, and have elucidated its roles in a wide range of disease and cancer contexts (*11-13*). Notably, ferroptosis is driven by peroxidation of membrane phospholipids enriched with polyunsaturated fatty acids (PUFAs), which is suppressed by glutathione peroxidase (GPX4) by catalyzing the reduction of phospholipid peroxides (*11, 14*). Therefore, the balance between monounsaturated fatty acid (MUFA) and PUFA phospholipids, together with GPX4 activity, plays a critical role in determining susceptibility to ferroptosis.

Here we investigate the molecular basis of castration response through *in vivo* analyses in genetically-engineered mice, supported by validation to human prostate tumors and organotypic culture assays. We show that ferroptosis represents the primary mechanism for luminal epithelial cell death in response to castration, and results from the combined inactivation of two distinct ferroptosis-suppressing pathways. We find that ferroptosis is normally suppressed in the hormonally-intact prostate by AR signaling within luminal epithelial cells through an intrinsic pathway mediated by the prostate-specific NKX3.1 homeodomain transcription factor and by extrinsic AR activity in the prostate stroma that regulates pleiotrophin (PTN), a secreted signaling factor that upregulates the mTORC1 pathway through stromal-epithelial interactions in epithelial cells. We demonstrate that inactivation of both the extrinsic and intrinsic AR signaling pathways is required to trigger prostate tissue regression and ferroptosis following androgen-deprivation.

## Results

### Defining key features of prostate regression

To understand castration response in the prostate, we initially sought to define the phenotypic and molecular features of prostate regression following castration. Our analyses have focused on alterations in luminal cells, since they represent the primary epithelial population and are known to undergo cell death following castration (*8, 15*). Notably, recent single-cell RNA-sequencing (scRNA-seq) analyses have revealed heterogeneity of luminal epithelial cells along the proximal-distal axis of each lobe in the mouse prostate (*16-20*), separating distal luminal (LumDist) cells, which correspond to traditional secretory cells, from a much smaller but distinct population of proximal luminal (LumP) cells that have progenitor properties (*16, 17*); analogous acinar and ductal (club) luminal epithelial populations have been described in the human prostate (*16-18, 20, 21*). Although both distal and proximal luminal cells express AR, most distal but not proximal luminal cells undergo cell death following androgen-deprivation (*18-20*).

In initial studies, we analyzed prostate tissues from wild-type C57BL/6 mice at three different time points following surgical castration (Fig. 1A), and observed progressive decreases in the size and weight of each prostate lobe at 3, 7, and 28 days post-castration (Fig. 1, B and C, and fig. S1, A and B), with 28 days considered to represent the fully regressed state(*15, 18*). We also observed decreased luminal cell size as measured in tissue sections, with a more pronounced reduction in distal luminal cells of each lobe (LumA, LumD, LumL, LumV) relative to proximal luminal cells (LumP) shared between all four lobes (Fig. 1D and fig. S1C). In addition, we found expression of proximal luminal markers such as PPP1R1B(*16*) in distal luminal cells starting at 7 days post castration (Cas d7) (Fig. 1B). This finding is consistent with scRNA-seq analyses showing a transcriptomic shift of surviving distal luminal cells towards proximal luminal states after castration (Fig. 1E), which has been previously reported (*18-20*).

**Fig. 1.**
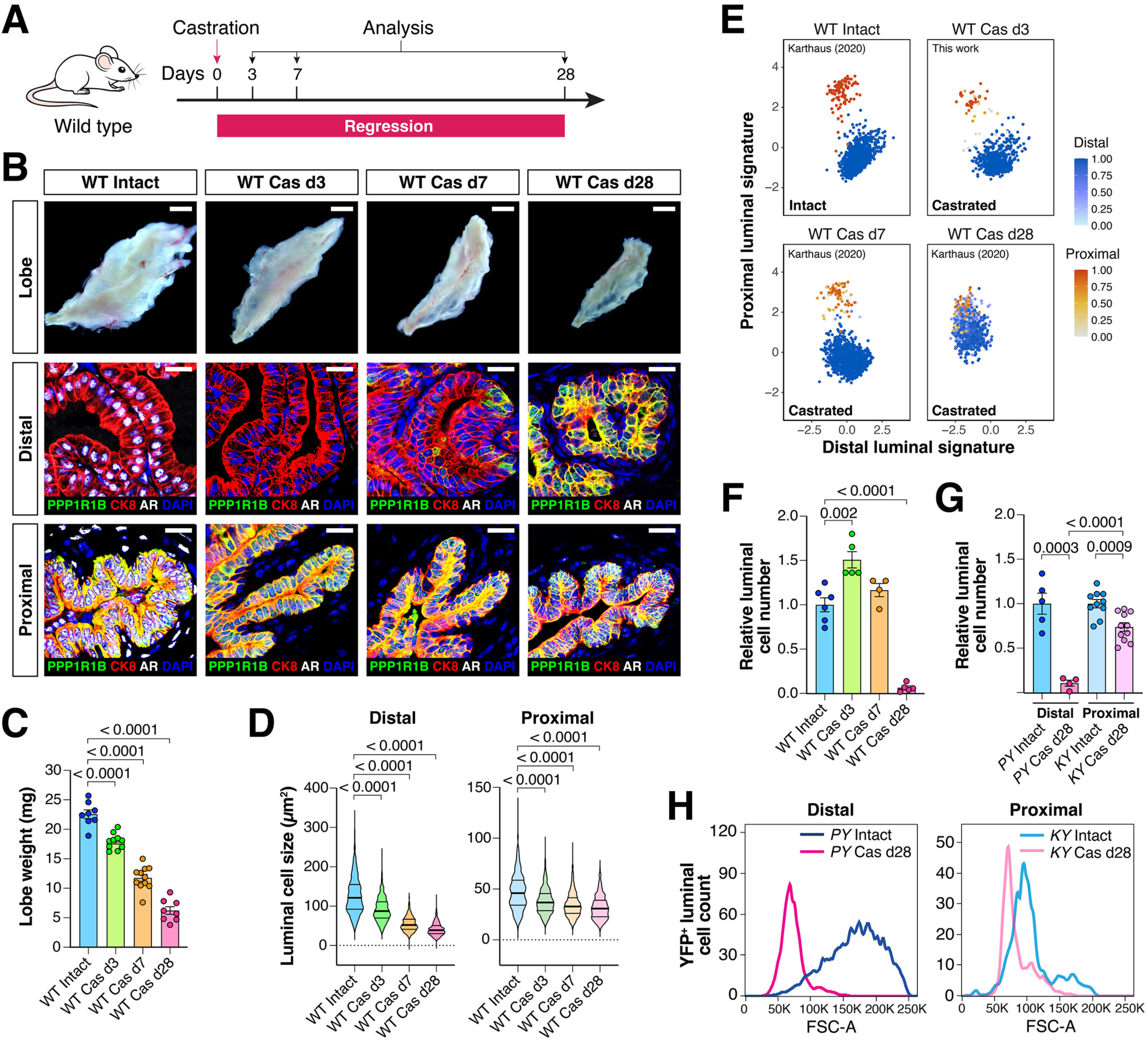
Defining features of prostate regression after castration. (**A**) Schematic of experimental analysis. (**B**) Dark-field images of whole-mount anterior prostate (AP) lobes and immunofluorescence staining of AP lobe sections from 10-12 week old adult wild-type (WT) mice that were hormonally intact or castrated for the indicated number of days. Scale bars for whole-mounts, 1 mm; for sections, 20 µm. (**C-D**) Prostate lobe weights (C) and distal and proximal luminal cell sizes (D) in AP lobes from intact and castrated WT mice. (**E**) Scatterplots of scRNA-seq data for AP luminal cells from intact or castrated mice, with each distal luminal (blue) or proximal luminal (red) cell assigned a distal (x-axis) and proximal luminal (y-axis) signature score. Color intensity corresponds to the probability strength of an XGBoost classifier assignment (color bar). (**F**) Luminal cell numbers in AP lobes from intact and castrated WT mice. (**G-H**) YFP-positive luminal cell number (G) and size (H) in AP lobes from intact and castrated tamoxifen-treated *PSA-CreER^T2^; Rosa26^LSL-YFP^*^/+^ mice (*PY*), which labels distal luminal cells, and *Krt7-CreER^T2^; Rosa26^LSL-YFP^*^/+^ mice (*KY*), which labels proximal luminal cells. All p-values were calculated using t-tests.

To identify changes in luminal cell number after castration, we used flow cytometry with calibrated counting beads to quantify luminal cell numbers in dissociated prostate lobes (fig. S1D). We found a transient increase in luminal cell number at Cas d3 followed by a minimal change at Cas d7, but a profound depletion of luminal cells (approximately 90%) by Cas d28 (Fig. 1F and fig. S1E). Notably, the increased number of luminal cells at Cas d3 is due to proliferation of distal luminal cells (fig. S2, A and D), consistent with the role of AR in suppressing luminal proliferation in the wild-type prostate epithelium (*22, 23*). To compare the response of distal versus proximal luminal cells to castration, we examined YFP-positive luminal cells at Cas d28 from tamoxifen-treated *PSA-CreER^T2^; Rosa26^LSL-YFP/+^* mice (PY) to label distal luminal cells and from *Krt7-CreER^T2^; Rosa26^LSL-YFP/+^* mice (KY) to label proximal luminal cells (fig. S1, F to H). We confirmed the increased expression of additional proximal luminal markers such as Sca-1 in distal luminal cells following castration (fig. S1I), and observed a significantly greater reduction in cell size and cell number in distal luminal cells compared to proximal luminal cells (Fig. 1, G and H). Together, these results show that we can define prostate regression through changes in lobe weight, luminal cell size, luminal transcriptomic state, and luminal cell number; however, these changes do not occur in a synchronized manner.

### Death of distal luminal cells after castration is mediated by ferroptosis

To evaluate cell death mechanisms following castration, we first examined the occurrence of apoptosis by cleaved caspase-3 and terminal deoxynucleotidyl transferase dUTP nick end-labeling (TUNEL) staining in prostate tissue sections from a time course after castration. We found that apoptotic distal luminal cells could be detected as early as Cas d1, with a peak of apoptosis in approximately 2% of distal luminal cells at Cas d3, declining to 0.5% or less between Cas d4 and Cas d28 (fig. S2, B to C and E to F). Importantly, the early and low rate of apoptosis together with the measured increase in cell proliferation at Cas d3 compared to the extensive loss of luminal cells at Cas d28 indicated that an alternative mechanism(s) promotes cell death after castration.

To identify key mechanisms involved in distal luminal cell death, we used genetically-engineered mouse models (GEMMs) to test requirements for cell death pathways *in vivo* following castration. For this purpose, we deleted key regulators of cell death pathways in prostate epithelial cells using the *Tmprss2^CreERT2^* inducible Cre driver (*24*) to determine whether cell death would be rescued after castration. We found that deletion of *Casp8*, which plays a crucial role in initiating apoptosis, particularly through the extrinsic (death receptor-mediated) pathway (*25*), or of *Gsdmd,* which is critical for pyroptosis (*26*), did not significantly alter cell death after castration (Fig. 2, A and B). In contrast, incomplete deletion of *Acsl4,* which encodes acyl-CoA synthetase long-chain family member 4, a key regulator of ferroptosis (*27*), partially rescued luminal cell death after castration (Fig. 2B). These findings are consistent with analyses of a published dataset (*28*) indicating that regressed prostate tissues *in vivo* are enriched for a signature of ferroptosis, but not for signatures of apoptosis, pyroptosis, or necroptosis (fig. S2G and Table S1). Notably, we found that treatment of wild-type mice with a potent inhibitor of lipid peroxidation, liproxstatin-1 (*29*), could dramatically suppress cell death following castration (Fig. 2, C to E).

**Fig. 2.**
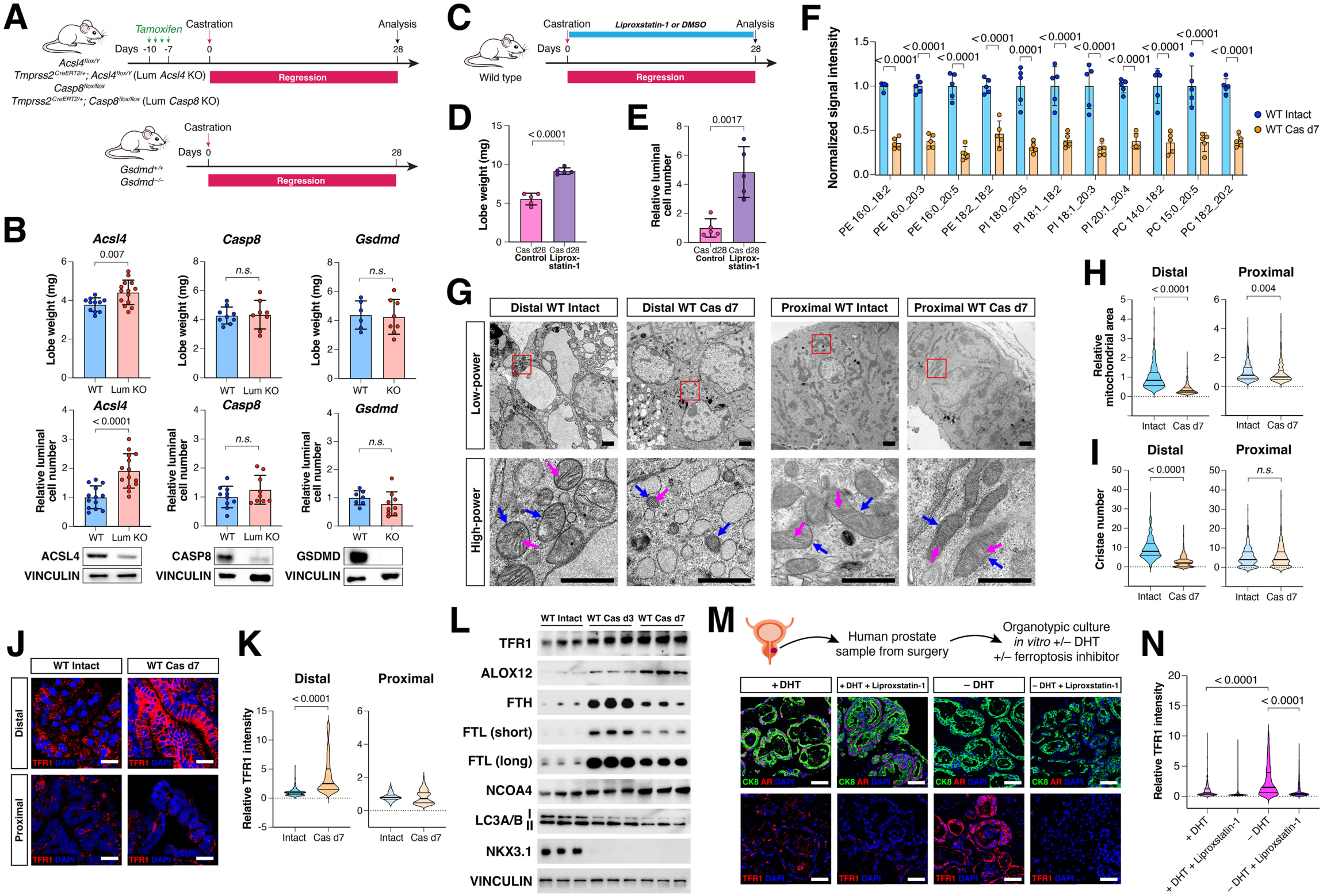
Luminal cell death after castration is mediated by ferroptosis. (**A**) Experimental strategy for testing the *in vivo* roles of ferroptosis, apoptosis, and pyroptosis in luminal cell death after castration. (**B**) Lobe weight and luminal cell number in AP lobes of control (WT) and mice with induced luminal deletion (Lum KO) for *Acsl4, Casp8,* or *Gsdmd* at 28 days after castration, with deletion efficiency confirmed by immunoblotting. (**C**) Strategy for blocking ferroptosis *in vivo* by daily administration of liproxstatin-1 or DMSO control by oral gavage. (**D-E**) Lobe weight and luminal cell number in control and liproxstatin-1 treated mice at Cas d28. (**F**) Representative poly-unsaturated fatty acid (PUFA)-phospholipids that are significantly downregulated in luminal cells from hormonally-intact and castrated wild-type mice at 7 days after castration (Cas d7), with each point corresponding to lipids extracted from combined luminal cells from 5 mice; see Methods for details. (**G**) Electron micrographs of distal and proximal luminal cells in AP lobes of hormonally-intact and Cas d7 wild-type mice at low-magnification (top) and high-magnification (bottom). Mitochondria (blue arrows) and cristae (pink arrows) are indicated. Scale bars, 1 µm. (**H-I**) Quantification of mitochondrial area (H) and cristae number (I). (**J-K**) Immunofluorescence staining of TFR1 in AP lobes from hormonally-intact or Cas d7 wild-type mice (J), with quantification of mean TFR1 fluorescence staining intensity in distal and proximal luminal cells (K). Scale bars, 20 µm. (**L**) Western blotting of ferroptosis-associated proteins in luminal cells from intact, Cas d3, and Cas d7 wild-type mice, with each lane corresponding to combined protein extracted from 5 mice. (**M-N**) Immunofluorescence staining of CK8, AR, and TFR1 in organotypic cultures of primary human prostate tumors grown for 4 days in the indicated conditions (M), with quantification of mean TFR1 staining intensity in CK8-positive tumor cells (N). Scale bars, 50 µm. All p-values were calculated using t-tests.

The detection of ferroptosis in tissues *in vivo* can be reliably ascertained by analysis of changes in lipid metabolism, alterations in mitochondrial morphology, and changes in expression of specific markers (*12*). Therefore, we performed lipidomic analyses of prostate tissues by liquid chromatography-mass spectrometry (LC-MS), which revealed substantial decreases in the profiles of polyunsaturated fatty acid (PUFA) phospholipids in luminal cells from prostates at Cas d7, but not at Cas d3 (Fig. 2F and fig. S3, A to C), consistent with lipid peroxidation and ferroptosis. We also examined mitochondrial morphology in regressed and intact prostate tissue sections by electron microscopy, which revealed decreases in mitochondrial size and number of cristae, a morphologic feature of ferroptosis (*30*), in distal luminal cells at Cas d7, but not proximal luminal cells (Fig. 2, G to I).

Next, we examined the expression of transferrin receptor (TFR1), a marker of ferroptosis (*31*), and observed a strong increase in TFR1 staining in distal luminal cells at Cas d7 (Fig. 2, J and K), consistent with Western blot analyses indicating a strong increase in TFR1 and arachidonate 12-lipoxygenase (ALOX12) (Fig. 2L). Using TFR1 staining to visualize the occurrence of ferroptosis from the same time course after castration as analyzed for apoptosis, we found upregulation of TFR1 expression in over 50% of distal luminal cells at Cas d7, which continued to be elevated at Cas d14 and Cas d28 (fig. S4, A to C). These findings were further confirmed by staining for 4-hydroxynonenal (4-HNE), a product of lipid peroxidation that is upregulated during ferroptosis (*32, 33*). We found that 4-HNE staining followed a similar time course as TFR1 expression, with increased levels in over 40% of distal luminal cells evident at Cas d7, d14, and d28 (fig. S4, D to F). Furthermore, we also examined the levels of labile Fe^2+^ by flow cytometry, and detected high levels in luminal cells at Cas d2 and Cas d3, consistent with an early event prior to the onset of ferroptosis (fig. S4, G and H). Taken together, these analyses indicate that the onset of ferroptosis during prostate regression occurs after the occurrence of apoptosis, and is the primary mediator of distal luminal cell death.

We next investigated the induction of ferritinophagy, which often plays a sensitizing role in ferroptosis induction (*34, 35*). Western blot analyses showed autophagy induction as detected by LC3 ratios in luminal cells at Cas d3 and d7 (Fig. 2L), together with degradation of ferritin heavy chain (FTH) and ferritin light chain (FTL) at Cas d7 (Fig. 2L). Notably, we also observed a transient increase in FTH and FTL expression along with upregulation of *Fth* and *Ftl* mRNA levels in luminal cells before Cas d7 (Fig. 2L and fig. S5D), analogous to early events after cells are treated with ferroptosis inducers *in vitro* (*34, 35*). Using LC3 reporter mice to examine autophagosomes (*36*), we performed a similar time course analysis of prostate regression, and found that LC3 puncta size was elevated at Cas d1 through Cas d7 (fig. S5, A to C), consistent with autophagy preceding ferroptosis. To further test the role of autophagy, we examined prostate tissues from mice with luminal-specific deletion of *Atg7* (Lum *Atg7* KO), an essential mediator of autophagy (fig. S5E). We found that these Lum *Atg7* KO mice displayed increased FTH and FTL levels, reduced TFR1 and ALOX12 expression, and partial rescue for cell size, cell number and lobe weight phenotypes, but displayed a less prominent transcriptomic shift after castration (fig. S5, F to M). Together, these genetic, lipidomic, ultrastructural, and biochemical findings indicate that castration induces prostate regression primarily through the mechanism of ferroptosis in distal luminal cells.

We validated our findings in mouse models by investigating ferroptosis in human prostate tissues following ADT. First, we observed TFR1 upregulation by immunostaining of a cohort of archival benign prostate tissues and tumors from patients at Columbia University Irving Medical Center who had undergone neoadjuvant ADT prior to prostatectomy or transurethral resection of prostate (TURP) (fig. S3, D and E and Table S2). Secondly, we examined a published scRNA-seq dataset containing castration-sensitive prostate cancers (CSPC) after ADT(*37*), and found enrichment of an expression signature for ferroptosis(*28*) in the castration-sensitive prostate cancers (CSPC) relative to the CRPC tumors (fig. S3F and Table S1). We also found that the ferroptosis signature was enriched in enzalutamide responders relative to non-responders in a bulk RNA-seq dataset of biopsies from metastatic CRPC patients(*38*) (fig. S3G). Finally, in analyses of organotypic cultures of human prostate tumor tissue (Table S3), we found that TFR1 expression was significantly increased in the absence of exogenous dihydrotestosterone (DHT) but was reduced by addition of the ferroptosis inhibitor liproxstatin-1 (Fig. 2, M and N). Taken together, these findings indicate that ferroptosis occurs in human prostate tissues following ADT.

### AR is required in both stroma and epithelium to suppress ferroptosis

To determine whether prostate regression results from loss of AR function in the epithelium or stroma, or both, we performed inducible deletion of the *Ar* gene *in vivo* (Fig. 3A). To delete *Ar* in the prostate epithelium (which we term Lum *Ar* KO), we used inducible *Tmprss2^CreERT2/+^; Ar^flox/Y^* mice. For efficient deletion of *Ar* in the stroma, we used the combination of two inducible stromal Cre drivers in *Vimentin-CreER^T2^; Col1a2-CreER^T2^; Ar^flox/Y^* mice (Str *Ar* KO), or all three Cre drivers to delete *Ar* in both the epithelium and stroma (Lum + Str *Ar* KO) (fig. S6, A and B). We found that Lum *Ar* KO resulted in decreased lobe weight, luminal cell size, and a transcriptomic shift of distal luminal cells, whereas Str *Ar* KO only led to decreased lobe weight and luminal cell size (Fig. 3, B to F and fig. S6C); however, neither resulted in increased cell death (Fig. 3E). In contrast, the combination of both luminal and stromal *Ar* deletion (Lum + Str *Ar* KO) resulted in loss of approximately 80% of distal luminal cells as well as substantial reductions in lobe weight, luminal cell size, and transcriptomic shift, similar to the effects of castration (Fig. 3, B to F).

**Fig. 3.**
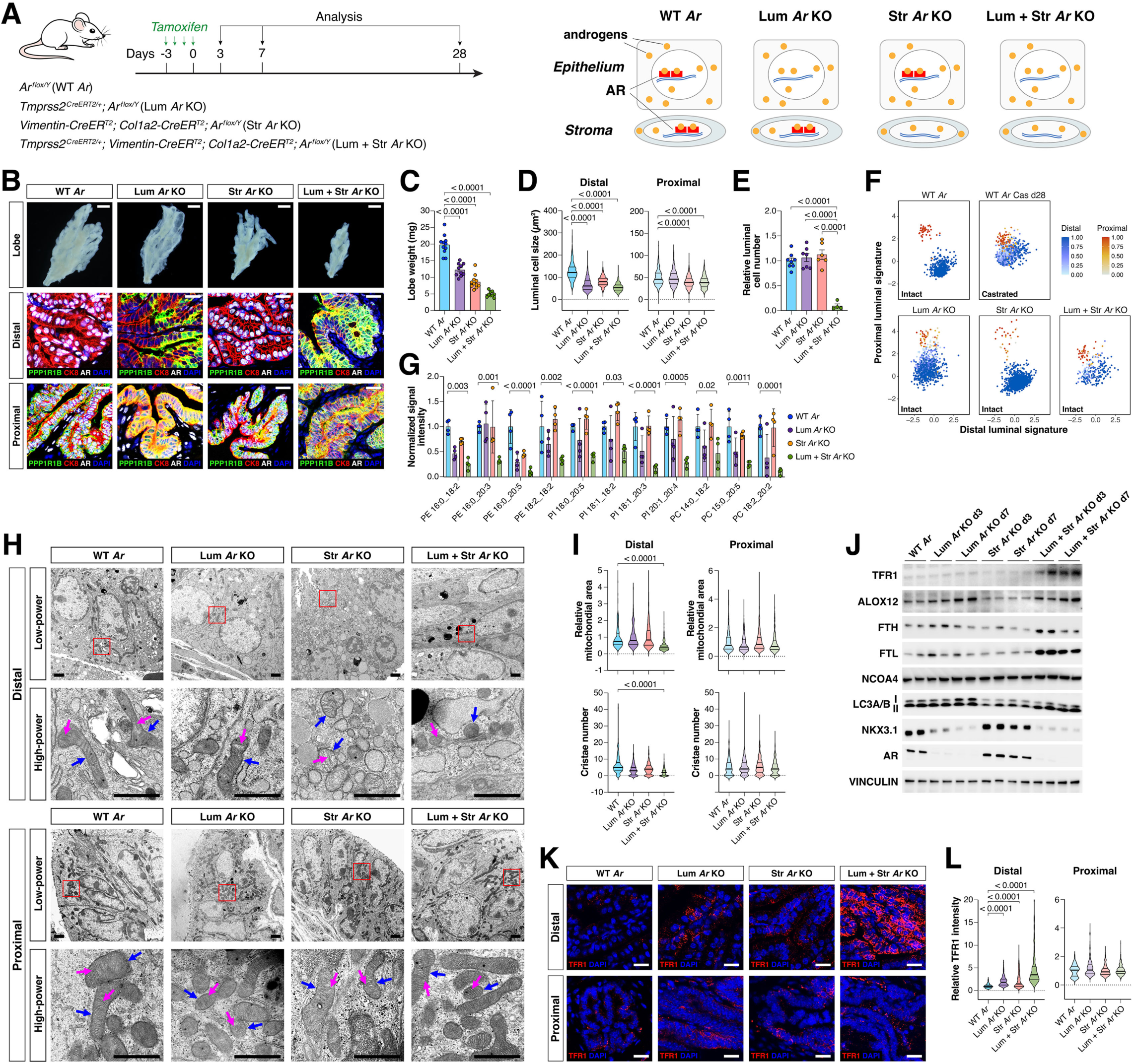
Androgen receptor has distinct roles in prostate epithelium and stroma. (**A**) Experimental strategy and mouse genotypes for inducible deletion of *Ar* in prostate luminal cells and/or stromal cells. (**B**) Dark-field images of AP lobes and immunofluorescence staining of AP sections from control mice and mice with induced deletion of *Ar* in luminal cells (Lum *Ar* KO), stromal cells (Str *Ar* KO), or both (Lum + Str *Ar* KO), at day 28 after tamoxifen induction. Scale bars for whole-mounts, 1 mm; for sections, 20 µm. (**C-E**) Lobe weight (C), luminal cell size (D), and luminal cell number (E) from mice of the indicated genotypes at day 28 after tamoxifen induction. (**F**) Scatterplots of scRNA-seq data showing transcriptomic changes in distal (blue) and proximal (red) luminal cells from control mice or mice with induced *Ar* deletion at day 28 after tamoxifen induction. (**G**) Abundance of representative PUFA-phospholipids in luminal cells from mice with the indicated genotypes at day 7 after tamoxifen induction, with each point corresponding to lipids extracted from combined luminal cells from 5 mice. Representative PUFA-phospholipids that were consistently altered in WT intact versus Cas d7 mice and Lum + Str *Ar* KO versus WT *Ar* mice at day 7 after tamoxifen induction are shown. (**H**) Electron micrographs of distal and proximal luminal cells in AP lobes from mice with the indicated genotypes at day 7 after tamoxifen induction, shown at low-magnification (top) and high-magnification (bottom). Mitochondria (blue arrows) and cristae (pink arrows) are indicated. Scale bars, 1 µm. (**I**) Mitochondrial size and cristae number from mice of the indicated genotypes. (**J**) Western blotting of ferroptosis-associated proteins in luminal cells from mice of the indicated genotypes at days 3 and 7 after tamoxifen induction, with each lane corresponding to combined protein extracted from 5 mice. (**K**) Immunofluorescence staining of transferrin receptor (TFR1) in AP lobes of mice with the indicated genotypes at 7 days after tamoxifen induction. Scale bars, 20 µm. (**L**) Mean fluorescence staining intensity of TFR1 in distal and proximal luminal cells. All p-values were calculated using t-tests.

To confirm that Lum + Str *Ar* KO promotes distal luminal cell death through ferroptosis, we performed lipidomic analyses to examine PUFA-phospholipid abundance. These analyses revealed significant reduction in PUFA-phospholipid levels in luminal cells from Lum + Str *Ar* KO prostates, but not Lum *Ar* KO or Str *Ar* KO prostates (Fig. 3G and fig. S6D). Consistent with these findings, ultrastructural analyses of mitochondrial morphology showed decreased mitochondrial size and cristae number in distal luminal cells but not proximal luminal cells from Lum + Str *Ar* KO prostates (Fig. 3, H and I). Similarly, Western blot analyses showed increased expression of TFR1 and ALOX12 in luminal cells from Lum + Str *Ar* KO prostates, but not in the other genotypes (Fig. 3J); in addition, we found increased TFR1 staining in distal luminal cells from Lum + Str *Ar* KO prostates (Fig. 3, K and L). Notably, we observed induction of autophagy in luminal cells from both Str *Ar* KO and Lum + Str *Ar* KO, suggesting that autophagy is regulated by stromal signals (Fig. 3J). Taken together, these findings indicate that loss of both extrinsic and intrinsic AR pathways mediate the effects of prostate regression after castration.

To elucidate the molecular mechanisms by which castration promotes ferroptosis, we performed pathway analyses comparing scRNA-seq data for prostate tissue at Cas d7 versus Intact(*18*), which showed that many of the pathways downregulated after castration are associated with lipid and fatty acid metabolism (fig. S7, A and B). Therefore, we focused on genes involved in the synthesis of MUFA- and PUFA-phospholipids, and found that genes related to MUFA-phospholipid synthesis including *Acsl3, Scd1, Fasn,* and *Mboat2* were downregulated at the mRNA and protein levels in luminal cells after castration (fig. S7, C and D); in contrast, expression of the PUFA-phospholipid associated gene *Fads2* showed significant upregulation (fig. S7D). Lipidomic analyses consistently showed a decrease in MUFA-phospholipids as early as Cas d3 (Fig. 4, A and B), prior to the reduction in PUFA-phospholipids at Cas d7 (Fig. 2F and fig. S3, A to C), suggesting that the downregulation of MUFA-phospholipid synthesis promotes ferroptosis induction. Notably, this downregulation was associated with decreased expression of MUFA-phospholipid biosynthesis genes in distal luminal cells following castration, and to a lesser extent in proximal luminal cells (fig. S7, E and F). We also observed a similar pattern of decreased expression of GPX4, a critical regulator of ferroptosis(*39*), in distal versus proximal luminal cells following castration (fig. S7, C to F).

**Fig. 4.**
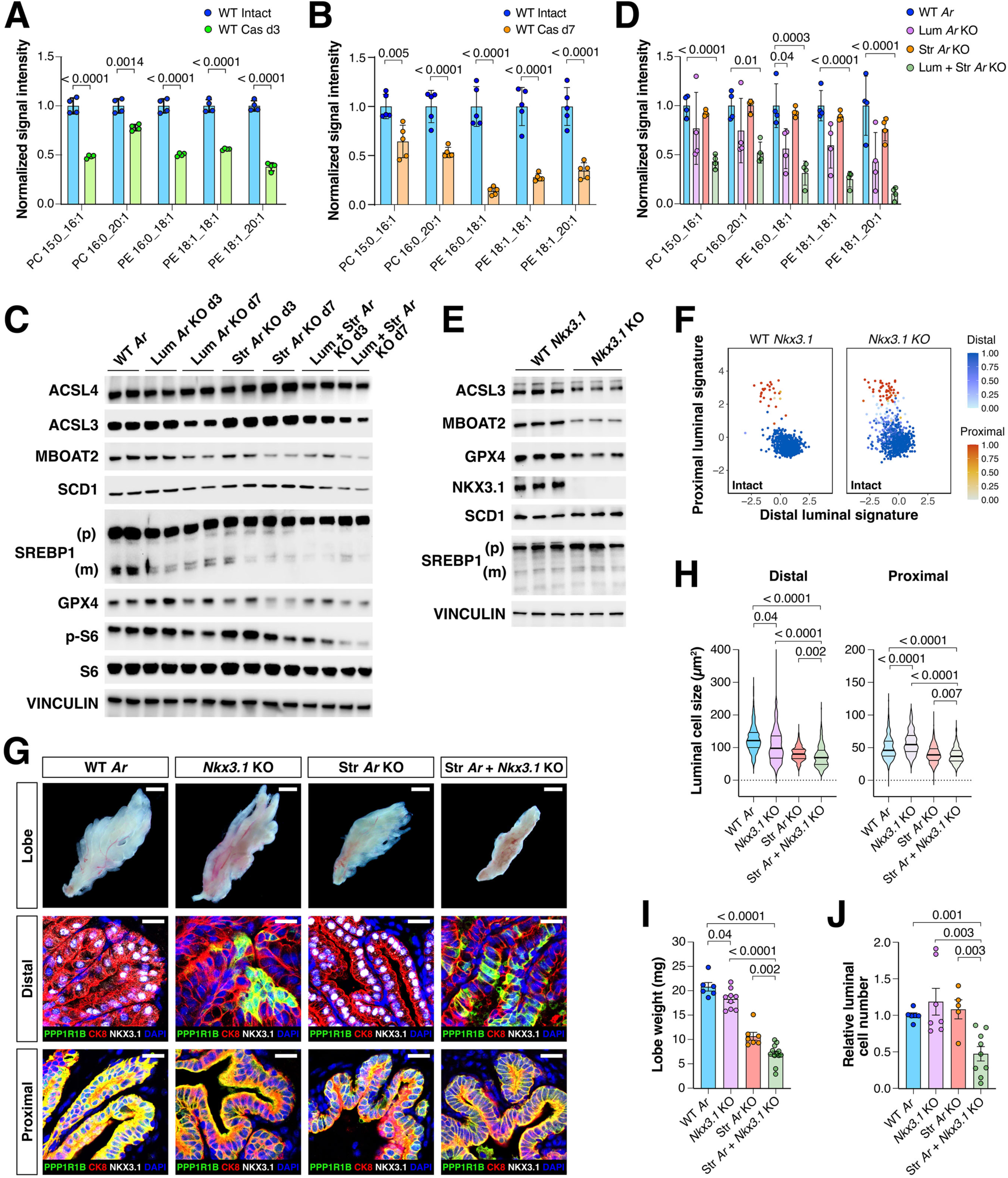
The intrinsic AR signaling pathway for castration response in luminal cells is mediated by *Nkx3.1*. (**A, B, D**) Abundance of representative MUFA-phospholipids in luminal cells from WT intact and Cas d3 mice (A), WT intact and Cas d7 mice (B), or from mice with induced deletion of *Ar* at day 7 after tamoxifen induction (D), with each dot corresponding to lipids extracted from combined luminal cells from 5 mice. Representative MUFA-phospholipids that were consistently altered in WT intact versus Cas d7 mice and Lum + Str *Ar* KO versus WT *Ar* mice at day 7 after tamoxifen induction are shown. (**C**) Western blotting of proteins involved in mTORC1 and MUFA/PUFA-phospholipid synthesis pathways in luminal cells from mice with induced deletion of *Ar* at day 3 and day 7 after tamoxifen induction, with each lane corresponding to combined protein extracted from 5 mice. (**E**) Western blotting of proteins associated with MUFA-phospholipid synthesis in luminal cells from WT and *Nkx3.1* mutant (*Nkx3.1* KO) mice, with each lane corresponding to combined protein extracted from 5 mice. (**F**) Scatterplots of scRNA-seq data showing transcriptomic changes in distal (blue) and proximal (red) luminal cells from control mice or *Nkx3.1* mutant mice. (**G**) Dark-field images of AP lobes and immunofluorescence staining of AP sections from control mice, and mice with *Nkx3.1* mutant (*Nkx3.1* KO), induced deletion of *Ar* in stromal cells (Str *Ar* KO), or both (Str *Ar* + *Nkx3.1* KO), at day 28 after tamoxifen induction. Scale bars for whole-mounts, 1 mm; for sections, 20 µm. (**H-J**) Luminal cell size (H), lobe weight (I), and luminal cell number (J) in mice of the indicated genotypes at day 28 after tamoxifen induction. All p-values were calculated using t-tests.

Given these findings, we examined expression of MUFA/PUFA-phospholipid biosynthesis genes and *Gpx4* in luminal cells from prostate tissues with luminal or stromal deletion of *Ar* or both. Our findings indicate that *Scd1, Fasn, Mboat2,* and *Gpx4* are co-regulated by both intrinsic and extrinsic AR pathways, whereas *Acsl3* and *Fads2* are solely regulated by the intrinsic pathway (Fig. 4C and fig. S7G). In addition, lipidomic analysis showed significantly decreased MUFA-phospholipid levels in Lum + Str *Ar* KO luminal cells, but not in Str *Ar* KO or Lum *Ar* KO luminal cells relative to wild-type luminal cells (Fig. 4D and fig. S6D). In combination, these findings show that the intrinsic and extrinsic AR pathways jointly regulate the expression of MUFA/PUFA-phospholipid biosynthesis genes as well as GPX4 to suppress ferroptosis in distal luminal cells.

### Nkx3.1 is a key mediator of the intrinsic castration-response pathway

Based on our prior work, we could readily identify a candidate regulator for the intrinsic pathway of castration-response, namely the homeodomain transcription factor NKX3.1. Notably, NKX3.1 is expressed in distal but not proximal luminal cells or the stroma (*16*) (fig. S7H), and its expression is regulated by AR (Fig. 3J) (*40, 41*); furthermore, NKX3.1 acts as a cofactor for AR binding to a subset of canonical AR targets (*42*). We found that expression of MUFA-phospholipid biosynthesis genes such as *Acsl3* and *Mboat2* as well as *Gpx4* was decreased, whereas the PUFA-phospholipid biosynthesis gene *Fads2* was upregulated in *Nkx3.1* null mutants (Fig. 4E and fig. S7, I and J). In addition, distal luminal cells in *Nkx3.1* mutants displayed a transcriptomic shift towards proximal states, resembling distal luminal cells from Lum *Ar* KO prostates (Fig. 4F). These findings are supported by our analyses of published RNA-seq and ChIP-seq data from mouse and human prostate (*43-46*), showing that the putative enhancers for *Acsl3, Mboat2, Gpx4* and *Fads2* are bound by NKX3.1 and AR, and that ADT reduces expression of these genes in human prostate tumors (fig. S7, K to M).

To test whether *Nkx3.1* acts in the intrinsic pathway to suppress ferroptosis, we asked whether *Nkx3.1* loss could effectively substitute for AR deletion in luminal cells in Str *Ar* + *Nkx3.1* KO mice (fig. S7I). After tamoxifen-induced deletion of *Ar* in the stroma for 7 days, the Str *Ar* + *Nkx3.1* mice displayed significantly increased TFR1 expression in distal luminal cells (fig. S7, N and O) as well as decreased expression of MUFA-phospholipid biosynthesis genes (fig. S7J). Furthermore, after tamoxifen-induced deletion of *Ar* in the stroma for 28 days, these mice displayed a significant decrease in lobe weight, luminal cell size, and increased luminal cell death (Fig. 4, G to J), similar to wild-type mice after castration as well as Lum + Str *Ar* KO mice. These findings indicate that *Nkx3.1* plays a central role in the intrinsic pathway of castration response.

### The extrinsic pathway is mediated by the AKT-mTORC1 pathway in luminal cells

To understand the properties of the extrinsic AR pathway in suppressing ferroptosis, we first sought to identify the signaling pathway(s) that receive stromal signals in distal luminal cells. Previous studies have implicated the AKT-mTORC1 pathway in regulating MUFA-phospholipid biosynthesis through the activity of the transcription factor SREBP1 (*47, 48*); in addition, mTORC1 regulates GPX4 expression at the level of translation (*49, 50*). Consistent with a key role of the AKT-mTORC1 pathway, we found reduced levels of phosphorylated AKT (p-AKT) and phosphorylated S6 protein (p-S6) together with decreased SREBP1 and GPX4 in luminal cells at Cas d7 (fig. S7C and fig. S8, A and B). Notably, the significant decrease in p-AKT, p-S6, and SREBP1 was observed in luminal cells of Str *Ar* KO and Str + Lum *Ar* KO prostates at 7 days after *Ar* deletion, but not in Lum *Ar* KO prostates (Fig. 4C and fig. S8, C and D). In addition, GPX4 protein levels were reduced in luminal cells in Str *Ar* KO prostates (Fig. 4C), but *Gpx4* mRNA levels were unaffected (fig. S7G), indicating that regulation of the extrinsic pathway occurs through translational control of GPX4.

To investigate whether the AKT-mTORC1 pathway acts downstream of stromal AR signaling, we performed luminal-specific inducible deletion of *Raptor,* which encodes a key component of mTORC1 (fig. S8E). We observed decreased p-S6, SREBP1, SCD1, MBOAT2 and GPX4 levels as well as ferritinophagy at 7 days after *Raptor* deletion in luminal cells (fig. S8F). Additionally, there was reduction of lobe weight and luminal cell size at 28 days, without any effects on transcriptomic state or luminal cell number (fig. S8, G to L). In mice with combined luminal deletion of *Raptor* and *Ar* (Lum *Raptor + Ar* KO), we found increased TFR1 expression in distal but not proximal prostate luminal cells at day 7 after tamoxifen treatment, consistent with induction of ferroptosis (fig. S8, M and N), as well as loss of luminal cell number at day 28 (fig. S8K), similar to wild-type mice after castration as well as Lum + Str *Ar* KO mice.

We further investigated the role of *Pten,* a tumor suppressor that negatively regulates AKT-mTORC1 signaling and plays a key role in prostate cancer progression as well as emergence of castration-resistance (*51, 52*). We found that luminal deletion of one allele of *Pten* (Lum *Pten* het KO) resulted in increased p-AKT levels and partially rescued expression of MUFA-phospholipid biosynthesis proteins, but had no detectable morphological phenotype in the prostate (fig. S9, A to E). In contrast to wild-type controls, however, Lum *Pten* het KO mice at Cas d28 displayed only a modest reduction in cell number (fig. S9, F to I), indicating that *Pten* heterozygosity partially suppresses ferroptosis induction. These findings are consistent with the known role of *Pten* mutants in promoting castration-resistance in prostate cancer.

### Pleiotrophin is the key signaling ligand for the extrinsic pathway

Next, we sought to identify the putative extrinsic signal(s) that act downstream of stromal AR to regulate activity of the AKT-mTORC1 pathway in luminal cells. Using bulk RNA-seq data from flow-sorted stromal cells, we focused on secreted signaling factors that exhibited significant changes (log_2_-fold change > 1.5, FDR corrected p-value < 0.05) at Cas d3 versus Intact (fig. S9J). Among the significantly downregulated stromal factors were Insulin-like Growth Factor 1 (IGF1) and Pleiotrophin (PTN, not to be confused with PTEN), which emerged as the top two candidates based on previous literature (*53-55*). Both *Igf1* and *Ptn* are highly expressed in prostate stromal cells, and their expression is strongly reduced after castration or stromal *Ar* deletion (fig. S9, K to N and fig. S10, A to D). In addition, to test whether IGF1 might act as the key stromal ligand, we used luminal-specific deletion of its receptor IGF1R to determine whether it would substitute for stromal AR loss (fig. S9O). However, we found that luminal deletion of *Igf1r* did not affect SREBP1 or p-S6 levels (fig. S9P), and had minimal phenotypes relative to Lum *Ar* KO prostates, with no significant effect on lobe weight and luminal cell number (fig. S9, Q to T). Thus, we concluded that IGF1 is unlikely to serve as a key stromal signal in the extrinsic AR pathway.

In parallel, we examined the phenotypes of inducible stromal-specific deletion of *Ptn* in Str *Ptn* KO mice (fig. S10E). Prostate tissues from Str *Ptn* KO mice displayed decreased AKT-mTORC1 pathway activity, as shown by reduced p-AKT and p-S6, decreased expression of GPX4 and MUFA-phospholipid biosynthesis genes, and induction of autophagy, similar to Str *Ar* KO prostates (Fig. 5A and fig. S10, F to I). However, the Str *Ptn* KO mice had a limited decrease in lobe weight and cell size, no alteration in transcriptomic states, and no effect on luminal cell number at 28 days after *Ptn* deletion (Fig. 5, B to E and fig. S10J). To test whether stromal deletion of *Ptn* would substitute for stromal *Ar* loss, we examined the phenotypes of Str *Ptn + Nkx3.1* KO mice, which combine stromal deletion of *Ptn* with null alleles for *Nkx3.1* (fig. S10, E and H). Notably, these mice test whether loss of this combination of putative intrinsic and extrinsic AR signaling pathway mediators could recapitulate prostate regression and ferroptosis without castration or *Ar* deletion. We found that the Str *Ptn + Nkx3.1* KO mice showed upregulation of TFR1 expression at day 7 following tamoxifen treatment (fig. S10, K and L), and expression of proximal markers in distal luminal cells, a strong transcriptomic shift, further decreases in lobe weight and luminal cell size, and most importantly a significant decrease in luminal cell number at day 28 after tamoxifen treatment (Fig. 5, B to E and fig. S10J). Most notably, we showed that treatment of wild-type mice with recombinant mouse PTN protein could significantly restore p-AKT and p-S6 levels, inhibit TFR1 induction at Cas d7 (fig. S10, M to P), and suppress the reduction of prostate lobe weight as well as the loss of luminal cells at Cas d28 (Fig. 5, F to H).

**Fig. 5.**
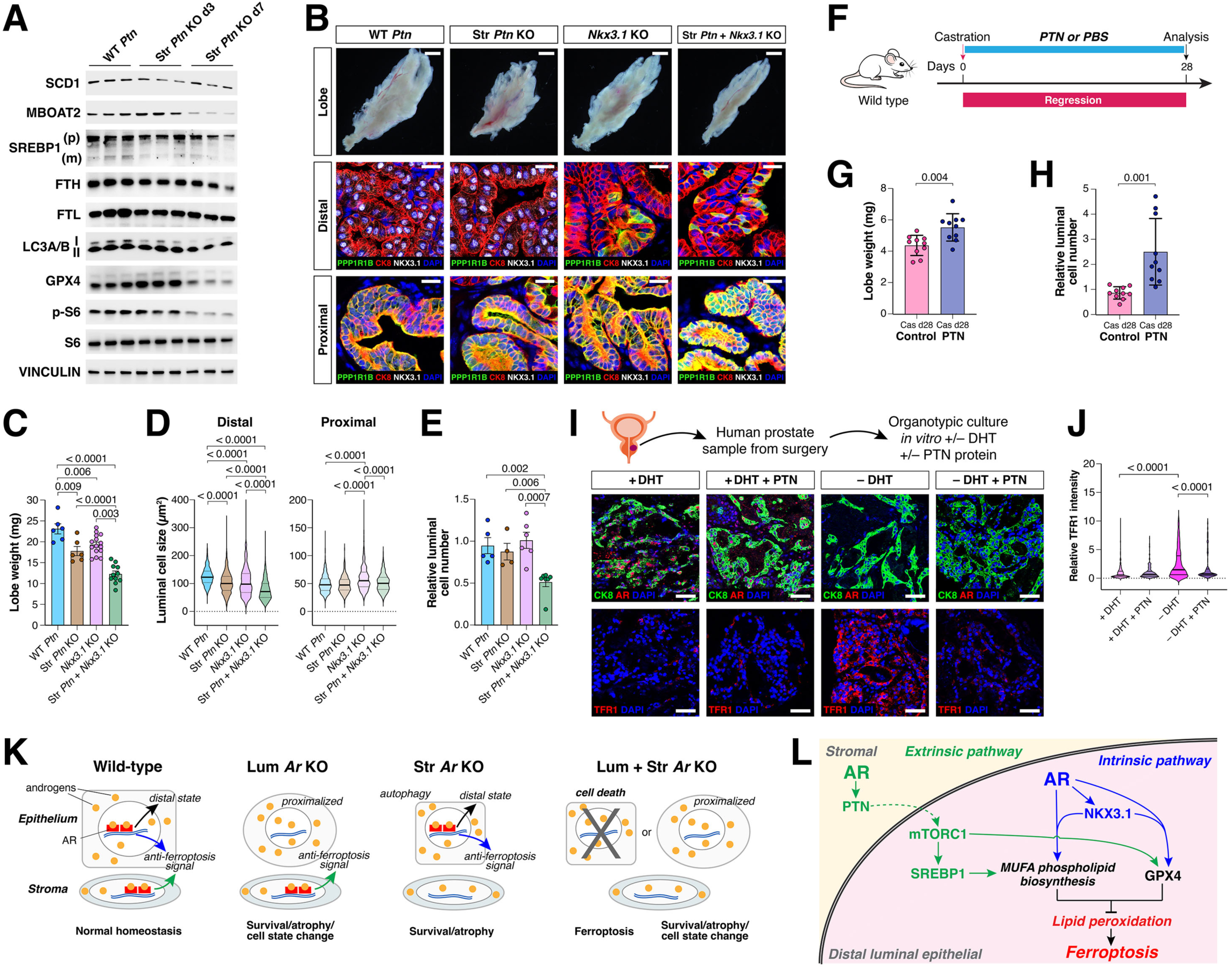
The extrinsic AR signaling pathway for castration response in luminal cells is mediated by a PTN-mTORC1 pathway. (**A**) Western blotting of proteins associated with mTORC1 and MUFA-phospholipid synthesis pathways in luminal cells from mice with induced stromal deletion of *Ptn* at day 3 and day 7 after tamoxifen induction. (**B**) Dark-field images and immunofluorescence staining of AP sections from control mice (WT *Ptn*), induced deletion of *Ptn* in stromal cells (Str *Ptn* KO), *Nkx3.1* mutant mice (*Nkx3.1* KO), or both (Str *Ptn* + *Nkx3.1* KO), at day 28 after tamoxifen induction. Scale bars for whole-mounts, 1 mm; for sections, 20 µm. (**C-E**) Lobe weight (C), luminal cell size (D), and luminal cell number (E) from mice of the indicated genotypes at day 28 after tamoxifen induction. (**F**) Strategy for blocking ferroptosis *in vivo* through daily administration of recombinant mouse PTN protein or PBS control via intraperitoneal injection. (**G-H**) Lobe weight and luminal cell number in control and PTN treated mice at Cas d28. (**I-J**) Immunofluorescence staining of CK8, AR, and TFR1 in organotypic cultures of primary human prostate tumors grown for 4 days under the specified conditions (I), together with quantification of mean TFR1 staining intensity in CK8-positive tumor cells (J). These cultures were performed simultaneously with those shown in Fig. 2M and 2N. Scale bars, 50 µm. All p-values were calculated using t-tests. (**K**) Model for epithelial and stromal AR roles in mediating castration-response. (**L**) Dual pathway mechanism for regulation of castration-response and ferroptosis.

Finally, to validate the key role of PTN in the extrinsic pathway, we examined whether exogenous PTN could suppress ferroptosis in organotypic cultures of human prostate tumor tissue (Table S3). We found that addition of recombinant human PTN protein could suppress expression of TFR1 in the absence of DHT (Fig. 5, I and J), resembling the effect of ferroptosis inhibitor liproxstatin-1 (Fig. 2, M and N), indicating that PTN can act as a potent extrinsic factor to suppress ferroptosis. Taken together, these findings suggest that PTN represents a key stromal signal that mediates signaling by the extrinsic AR pathway that cooperates with the NKX3.1-mediated intrinsic AR pathway to suppress ferroptosis in distal luminal cells.

## Discussion

AR signaling has been known for many decades to be of central importance for prostate organogenesis, adult homeostasis, and tumorigenesis, yet the molecular details of how loss of AR signaling results in luminal epithelial cell death have not been previously understood. We have used *in vivo* analyses in genetically-engineered mice together with validation in human prostate cancer to demonstrate that ferroptosis is the primary mechanism of luminal epithelial cell death after castration and is regulated through the integration of intrinsic and extrinsic AR signaling pathways. However, these intrinsic and extrinsic AR signaling pathways play distinct roles in castration response, as they are not equally important for each feature of tissue regression and ferroptosis (Fig. 5K).

Although induction of ferroptosis requires inactivation of intrinsic and extrinsic pathways, the intrinsic pathway plays a more fundamental role in regulating lipid metabolism (fig. S6D), and is exclusively involved in regulating the transcriptomic shift in distal luminal cell states (Fig. 3F). Moreover, the expression of NKX3.1 in distal luminal cells (fig. S7H) (*16*) and its role in regulating expression of *Gpx4* and MUFA/PUFA phospholipid biosynthesis may explain why distal luminal cells are more susceptible to ferroptosis than proximal luminal cells, and why prostate cancer patients with low NKX3.1 expression are more likely to benefit from treatment with 5-alpha-reductase inhibitors (*56*). In contrast, epithelial autophagy after castration is primarily mediated by the extrinsic pathway, as shown by our analyses of Str *Ar* KO mice (Fig. 3J).

Interestingly, the importance of stromal signaling in mediating epithelial cell death after castration was first recognized by Cunha using classical tissue recombination approaches to study stromal-epithelial interactions (*10*). We have identified the secreted ligand PTN as the stromal secreted factor that mediates this process through upregulation of mTORC1 activity to suppress ferroptosis induction and maintain adult homeostasis. In addition, PTN has recently been identified as a serum-based biomarker for pro-metastatic prostate cancer (*57*), and elevated serum levels of PTN are associated with a greater risk of lethal prostate cancer in African American men (*58*), suggesting that these associations are due to an increased propensity to develop castration-resistance through suppression of ferroptosis. Notably, the integration of dual pathways provides a fail-safe mechanism to prevent widespread prostate cell death in contexts of tissue damage that do not involve decreased AR signaling (Fig. 5L).

The mechanisms of prostate regression and regeneration following androgen deprivation and restoration likely represent an evolutionarily conserved feature of secondary sexual dimorphism in male mammals that undergo seasonal mating cycles, unlike mice and humans (*59*). For example, animals such as raccoons, white-tailed deer, and ground squirrels exhibit significant differences in the size of the prostate and other secondary sexual tissues between mating season and the rest of the year, when these tissues are highly regressed (*60-62*). This evolutionary conservation provides a strong rationale for the existence of two distinct regulatory pathways. The extrinsic pathway may be required to regulate tissue size through coordination of regression between stromal and epithelial cells, as has been hypothesized for organogenesis (*1, 3*); such a role may explain why the extrinsic pathway has a primary role in governing autophagy and cell size. Conversely, the intrinsic pathway is fundamental for maintenance of surviving distal luminal cells after castration, which are essential for regeneration of distal luminal cells after androgen-mediated regeneration. The cell state change due to the transcriptomic shift confers castration-resistance upon these residual distal luminal cells, which can then survive and repopulate the prostate during androgen-mediated regeneration (*17, 18*). Given this essential role in distal luminal survival, this function of the transcriptomic shift would logically be controlled by the intrinsic pathway.

It has long been appreciated that fatty acid biosynthesis is particularly important for prostate cancer, which requires *de novo* lipogenesis (*63, 64*). Notably, upregulation of FASN and MBOAT2 occurs in prostate tumor progression, and reprogrammed lipid metabolism is critical for prostate tumorigenesis (*65, 66*). Furthermore, MBOAT2 has been identified as an AR-regulated suppressor of ferroptosis through remodeling of phosphatidylethanolamine (PE) lipids to reduce oxidizable PUFA-phospholipids (*66*). Consequently, the central role of the PTEN-AKT-mTOR-SREBP1 pathway in castration response and resistance is due to the key role of SREBP1 in regulating fatty acid biosynthesis (*47, 67*). Somewhat unexpectedly, this pathway represents the downstream component of the extrinsic pathway, which is likely due to its relatively indirect role in lipogenesis. In contrast, we find that the intrinsic pathway plays a direct role in regulating the expression of key fatty acid biosynthesis genes, including *Mboat2*, *Scd1, Acsl3*, and *Fads2*. Moreover, expression of the key anti-ferroptotic protein GPX4 is regulated by both extrinsic and intrinsic pathways, at the levels of translation and transcription, respectively (Fig. 5L).

Since ferroptosis represents the physiological mechanism for epithelial cell death after androgen-deprivation of the normal prostate as well as hormone-sensitive prostate tumors *in vivo,* our findings suggest that inhibition of anti-ferroptotic signals, such as dual suppression of MUFA-phospholipid synthesis and GPX4, might serve as a replacement or supplementation for ADT in treatment of hormone-sensitive prostate tumors to reduce tumor recurrence and/or prevent androgen-deprivation induced lineage plasticity (*6, 68, 69*). Additionally, our findings suggest that promotion of ferroptosis combined with ADT may also enhance the response to ADT in treatment-naïve tumors and re-activate cell death responses for effective treatment of CRPC. Furthermore, since ferroptosis can propagate across tissues *in vivo* (*33, 70*), ferroptosis induction may represent an effective therapy even for heterogeneous cell populations within tumors.

Finally, since dietary components that influence oxidative states and lipid metabolism can modulate ferroptosis (*71*), our findings provide insights into the relationship between dietary patterns and prostate cancer incidence (*72*). Notably, our findings may help explain the increased risk of prostate cancer in the SELECT trial for healthy men with dietary supplementation with vitamin E (*73*), a potent anti-oxidant that can suppress ferroptosis (*71*). Therefore, the potential efficacy of combined pharmacological and dietary interventions for treatment of prostate cancer through ferroptosis induction represents a promising avenue for future translational studies.

## Materials and Methods

### Mice

Wild-type C57BL/6 male mice were purchased from Taconic Biosciences (C57BL/6NTac) at the age of 10-12 weeks. The mouse strains *PSA-CreER^T2^, Krt7-CreER^T2^, Tmprss2^CreERT2^*, *Vimentin-CreER^T2^, Acsl4^flox/flox^*, *Nkx3*.*1^-/-^, Atg7^flox/flox^*, *Ar^flox/Y^* and *Ptn^flox/flox^* mice have been previously described (*24, 74-81*); the *Ptn^flox/flox^* mice were provided by the Chute lab (Cedars Sinai Medical Center). The mouse strains *Col1a2-CreER^T2^* (Jax #029567), *Casp8^flox/flox^* (Jax #044034), *Gsdmd^-/-^*(Jax #032410), *Raptor^flox/flox^* (Jax #013188), *Igf1r^flox/flox^* (Jax #012251), *Pten^flox/flox^* (Jax #006440), *CAG-RFP-EGFP-LC3* (Jax #027139), and *Rosa26^LSL-YFP/+^* (Jax #006148) were obtained from The Jackson Laboratory. All mouse strains were maintained on a mixed C57BL/6-129Sv background. Tamoxifen induction was performed at 3 months of age by oral delivery of tamoxifen (Sigma-Aldrich T5648) at 150 mg/kg daily for 4 consecutive days. Liproxstatin-1 (Selleckchem S7699) was sequentially dissolved in 2% DMSO, 40% PEG300, 2% Tween 80, and 56% ddH_2_O, and was administered orally to castrated adult mice at a dosage of 30 mg/kg daily for 28 days. Pleiotrophin (PTN, R&D Systems 6580-PL-050) was dissolved in PBS and was injected intraperitoneally to castrated adult mice at a dosage of 300 ug/kg daily for 7 days or 28 days. All procedures followed protocols approved by the Institutional Animal Care and Use Committee (IACUC) at Columbia University Irving Medical Center.

### Dissociation of mouse prostate tissue

Prostate tissues were minced with scissors and incubated in papain (20 units/ml) with 0.1 mg/ml DNase I (Worthington LK002147) at 37°C with gentle agitation. After 60 min, the samples were gently triturated 30 times, and then incubated for an additional 45 min in papain. Samples were gently triturated again 30 times, followed by quenching of the enzyme using 1 mg/ml ovomucoid/bovine serum albumin solution with 0.1 mg/ml DNase I (Worthington LK002147) at 37°C for 5 min. Cells were washed with PBS-EDTA with 0.1 mg/ml DNase I (Worthington LS002145), followed by digestion in TrypLE Express (Invitrogen 12605-036) for 3-5 min at 37°C with gentle agitation. After spinning down for 5 min at 4°C, the cell pellet was resuspended in 1x PBS with 2% FBS and 0.1 mg/ml DNase I. Cells were passed through a 70 mm strainer (pluriSelect 43-10070-70) and a 20 mm strainer (pluriSelect 43-10020-70), washed in 1x PBS, and resuspended in appropriate buffers for downstream analyses.

### Quantification of luminal cell number

Prostate anterior lobes were micro-dissected, and the stroma was removed under the dissection microscope before performing dissociation. Dissociated prostate cells were stained with BV421-Sca1, APC-Epcam, and PE-CD66a antibodies in 100 μl HBSS with 2% FBS on ice for 30 min. 50 μl CountBright™ Absolute Counting Beads (ThermoFisher Scientific C36950) were added before loading on the BD LSRFortessa™ Cell Analyzer. Epcam^+^CD66a^high^ luminal cells were defined as described(*16*). For lineage-traced prostate epithelial cells, the Epcam^+^CD66a^high^EYFP^+^ cells from *PSA-CreER^T2^;Rosa26^LSL-YFP/+^* mice and *Krt7-CreER^T2^;Rosa26^LSL-YFP/+^* mice were defined as distal luminal cells and proximal luminal cells, respectively. The number of luminal cells was calculated by dividing luminal cell counts by counting bead counts and then multiplying by the quantity of counting beads added. Catalog numbers and dilutions for all antibodies are available in Table S4.

### Western blotting

Dissociated prostate cells were stained with eFluor450-CD45, eFluor450-Ter119, eFluor450-CD31, PE/Vio770-Sca1, APC-Epcam, and PE-CD66a antibodies in 100 μl of HBSS with 2% FBS on ice for 30 min. When sorting with a BD Influx cell sorter, DAPI (Sigma-Aldrich D9542, 10ug/ml) was added to remove dead cells. Sorted luminal cells were defined as EpCAM^+^CD66a^high^Lineage^-^ (CD45, CD31, and Ter119) and DAPI-negative cells. Cells were centrifuged at 300 g for 5 min at 4°C, and the cell pellet was resuspended in RIPA buffer (Sigma R0278) with a protease inhibitor cocktail (Sigma-Aldrich P8340) and phosphatase inhibitor cocktail (Sigma-Aldrich P0001) and incubated on ice for 30 min. After incubation, samples were centrifuged at 12,000 g for 15 minutes, and the supernatants were collected. Protein concentration was quantified using the DC Protein Assay Kit (Bio-Rad 5000112) before adding 4x Laemmli Sample Buffer (Bio-Rad 1610747), and the protein lysates were boiled for 5 minutes on a heating block and stored at –80°C. 20 μg protein was resolved on 4–15% Mini-PROTEAN TGX™ Precast Protein Gels (Bio-Rad 4561086), transferred to Immun-Blot PVDF Membrane (Bio-Rad 1620174), blocked in 5% non-fat milk in PBS with 0.1% Tween-20, incubated with primary antibodies overnight at 4°C, and further incubated with horseradish peroxidase (HRP)-linked anti-rabbit IgG or anti-mouse IgG after washing. Blots were imaged using a ChemiDoc MP imaging system (Bio-Rad 17001402). Catalog numbers and dilutions for all antibodies are available in Table S4.

### Quantitative real-time PCR

Epcam^+^CD66a^high^Sca1^low^Lineage^-^(CD45, CD31, and Ter119), DAPI-negative distal luminal cells, Epcam^+^CD66a^high^Sca1^high^Lineage^-^(CD45, CD31, and Ter119) DAPI-negative proximal luminal cells, and Epcam^-^Cd66a^-^Lineage^-^(CD45, CD31, and Ter119) DAPI-negative stromal cells were sorted from prostates of mice with specific genotypes. Total RNA was extracted using the Direct-zol™ RNA MicroPrep kit (Zymo Research R2060) according to the manufacturer’s protocol. cDNA was synthesized using PrimeScript™ RT Master Mix (Takara Bio RR036A), and real-time PCR was conducted with PowerUp™ SYBR™ Green Master Mix (Invitrogen A25742) on the Applied Biosystems 7500 FAST Real-Time PCR machine. Primers are detailed in Table S5.

### Bulk RNA-sequencing of prostate stromal cells

Epcam^-^Cd66a^-^Lineage^-^ (CD45, CD31, and Ter119), DAPI-negative stromal cells were sorted from dissociated hormonally-intact and Cas d3 adult wild-type mouse prostates. Sorted prostate stromal cells (∼1x10^5^ cells/mouse) were centrifuged at 300 g for 5 min at 4°C and resuspended in 100 μl TRIzol reagent (ThermoFisher Scientific 15-596-018). RNA was extracted using the Zymo Direct-zol RNA microprep kit (Zymo Research R2060), and the quality of extracted RNA was assessed on an Agilent Bioanalyzer® RNA 6000 Nano/Pico Chip. High-quality RNA with a RIN score > 7 was used for RNA library preparation using the NEBNext Ultra II Directional RNA Library Prep Kit for Illumina (NEB E7760). Libraries were sequenced on an Illumina HiSeq 2x150 bp platform (Azenta Life Sciences).

### Lipidomic analysis

Epcam^+^Cd66a^high^Lineage^-^ (CD45, CD31, and Ter119), DAPI-negative luminal cells were sorted from dissociated mouse prostates with the indicated treatments. The cell pellet was flash-frozen in liquid nitrogen and stored at -80°C until lipid extraction. For lipid extraction, 250 μl cold methanol containing 0.01% w/v butylated hydroxytoluene (BHT) was added to cell pellets. Cells were lysed by an ultrasonic homogenizer for 5 cycles. 850 μl of cold methyl tert-butyl ether was added to the lysate and vortexed at the highest speed for 30 seconds and then shaken on ice for at least 2 hr to enhance the lipid extraction. Then, 200 μl of MS-grade cold water was added to each vial, vortexed for 30 seconds, and incubated on ice for 10 min. The mixture was then centrifuged at 4,000 rpm for 20 min at 4°C to achieve phase separation. The organic layer containing lipids was collected in a separate glass vial and dried under a gentle stream of nitrogen. The final LC-MS sample was prepared by dissolving dried lipid in 4:3:1 (v/v/v) isopropanol (IPA):acetonitrile (ACN):water with 0.001% w/v BHT.

Samples were analyzed using an Acquity UPLC I-class PLUS interfaced with a Synapt G2-Si Mass spectrometer (Waters Corp.). Chromatographic separation was performed with a 20 min gradient elution profile on a Waters Acquity CSH C18 column (1.7 μm, 2.1 mm × 100 mm). Mobile phase: (A) water: ACN (40:60; v/v) and (B) water: ACN: IPA (5:10:85; v/v/v), both containing 10 mM ammonium acetate and 0.1% acetic acid. The following linear gradient at 400 µL/min flow rate with a column temperature at 55°C was used: 0-2 min: 60% B, 2-2.3 min: 75% B, 2.3-10 min: 90% B, 10-17 min: 100% B, 17-20 min: 40% B. The Synapt G2-Si mass spectrometer was equipped with a LockSpray ion source and was operated in both ESI ionization modes over the mass range of 50-1600 m/z. Source voltages were set to ±2 kV, 30 V, and 5 V for capillary, sampling, and extraction cones, respectively, and the temperature was set to 120°C for the source and 500°C for sample desolvation. Gas flow rates were set at 750 L/hr and 50 L/hr for the desolvation gas and cone gas, respectively. Fragment ion spectra were generated using enhanced data-independent ion mobility (HDMSE) acquisition mode where data from mobility separation ions are collected in two channels, with either low collision energy applied at 4 V or with an elevated collision energy ramp from 25 to 60 V for precursor and fragment ions, respectively. Nitrogen as the drift gas was held at a flow rate of 90 ml/min in the IMS cell with a wave velocity of 600 m/s and a wave height of 40 V.

Heatmaps of significantly altered lipid species were generated using a fold-change threshold of 1.5 and an FDR-corrected p-value of less than 0.05 between the specified groups. Representative PUFA- and MUFA-phospholipids that display consistent differences in levels in WT Cas d7 versus WT Intact and Lum + Str *Ar* KO versus WT *Ar* at day 7 after tamoxifen treatment are shown.

### Immunofluorescence staining

Immunofluorescence was performed on 5 μm formalin-fixed paraffin-embedded (FFPE) sections. FFPE sections were de-paraffinized and rehydrated, followed by heat-induced antigen retrieval using a citrate buffer (VectorLabs, pH 6.0) or Tris-EDTA buffer (VectorLabs, pH 9.0). Sections were blocked in 5% goat serum (ThermoFisher Scientific 50-062Z) in PBS at room temperature for 30 min. Incubation of primary antibodies diluted in 5% goat serum was performed at 4°C overnight. AlexaFluor-conjugated secondary antibodies raised in goats were used for signal detection. Slides were counterstained with 1 μg/ml 4’,6-Diamidino-2-Phenylindole (DAPI, ThermoFisher Scientific D1306) and mounted with coverslips using Prolong^TM^ Gold Antifade (ThermoFisher Scientific P36930). The Tyramide Signal Amplification (TSA) system was used and performed according to the manufacturer’s protocol (Akoya Biosciences NEL704A001KT) to detect immunofluorescence signals from TFR1, 4-HNE, p-AKT, and p-S6 staining. Images were captured using a Nikon Eclipse 50i microscope or on the Leica STELLARIS Confocal Platform. Catalog numbers and dilutions for all antibodies are provided in Table S4.

### Quantitative image-based cytometry

Images were acquired using a Nikon Eclipse 50i microscope equipped with a PL Apo 60X/0.95 numerical aperture (NA) objective. Acquisition times for the different channels were adjusted to obtain images in non-saturating conditions for all the treatments analyzed within the experiment. Depending on tissue size, 30 to 40 images were acquired, containing a total of 4,500 to 8,000 cells from 3 to 5 mice per condition. Using ImageJ, the CK8 channel was used to outline regions of interest (ROIs) that identified each luminal cell as an individual object. These ROIs were then applied to the TFR1 or 4HNE1 channel to quantify pixel intensities for each individual cell/object. After pixel quantification, the desired quantified values for each cell/object (mean intensity and area) were extracted and exported to Spotfire software (Revvity Signals). Spotfire was used to quantify percentages in cell populations and to generate all color-coded scatter diagrams.

### Single-cell RNA-sequencing

Dissociated cells were washed twice in 1x HBBS with 2% FBS, passed twice through 20 mm strainers (pluriSelect 4310020-70), and incubated on ice with hash-tag antibody (BioLegend, TotalSeq Panel B, B0301-B0305, 1 µl/test). After 30 min staining, cells were centrifuged at 300 g for 5 min at 4°C and washed three times in 1x PBS with 0.04% BSA. Cells were counted using a Countess II FL Automated Cell Counter (ThermoFisher Scientific). If the viability of the samples was greater than 80% and the single-cell fraction was greater than 95%, the hashtag-labeled cells were combined and resuspended in 1x PBS with 0.04% BSA at approximately 1 x 10^6^ cells/ml. Samples were submitted to the Columbia JP Sulzberger Genome Center for single-cell RNA sequencing on the 10X Genomics Chromium platform. Libraries were generated using the Chromium Single Cell 3’ Reagent Kit v2 for library construction. Samples were sequenced on a NovaSeq 6000 or NovaSeqX sequencer.

### Labile iron measurements

Dissociated prostate cells were stained using BioTracker FarRed Labile Iron dye (Sigma-Aldrich, SCT037, 5 μM) at 37 °C for 1 hr, washed three times with HBSS, and further stained with eFluor450-CD45, eFluor450-Ter119, eFluor450-CD31, APC/Cyanine7-EpCAM, and PE-CD66a antibodies in 100 μl 1x HBSS with 2% FBS on ice for 30 min. The stained cells were then spun down and resuspended in HBSS containing 10 μg/ml DAPI to exclude dead cells. Luminal cells were identified as Epcam^+^CD66a^high^Lineage^-^ (CD45, CD31, and Ter119) cells, and the APC channel was used to determine labile iron level in the gated luminal cells. Data were collected and analyzed using a BD LSRFortessa Cell Analyzer and FlowJo software, respectively.

### Human samples and organotypic culture assay

Human prostate specimens were obtained following protocols approved by the Human Research Protection Office and the Institutional Review Board (IRB) at Columbia University Irving Medical Center, with written informed consent from the patients. For immunofluorescence analyses, we generated a cohort of archival benign prostate tissues and tumors from patients at Columbia University Irving Medical Center who had undergone neoadjuvant ADT prior to prostatectomy or transurethral resection of prostate (TURP) (Table S2).

For organotypic culture, fresh prostate tissues were obtained immediately following surgical removal, and only tissues that were not needed for clinical diagnosis were used. Tissues were sectioned into 300 μm slices using a vibratome (Precisionary Instruments VF-510-0Z) and transferred to cell culture inserts (Falcon 08-771-12) with medium added to the plate. The medium contained 50% Keratinocyte-Serum Free Media (ThermoFisher Scientific 17005042), 50% Medium 199 (Sigma-Aldrich, M4530), 10 μM Rock Inhibitor Y-27632 (MedChemExpress HY-10071), and 100 μg/ml Primocin (InvivoGen ant-pm-2). Depending upon the experiment, we also added 10^-9^ M dihydrotestosterone (DHT) (Cayman Chemical 15874), 200 nM Liproxstatin-1 (Selleckchem S7699), and/or 300 ng/mL Pleiotrophin (PTN, R&D Systems 252-PL-250). Tissue slices were cultured for 4 days, collected and fixed in 10% formalin overnight, and processed to create paraffin blocks for downstream analysis. Information about the human patient samples for organotypic culture is provided in Table S3.

### Electron microscopy

Prostate AP lobes were prepared for pre-embedding for electron microscopy as described (*82, 83*). Experimental procedures were approved by the Institutional Animal Care and Use Committee of Weill Cornell Medicine. After deep sodium pentobarbital (150 mg/kg, *i.p.*) anesthesia, mouse prostate AP lobes were fixed with an aortic arch perfusion with ∼5 ml of 2% heparin in normal saline followed by 30 ml 2% paraformaldehyde (PFA) and 3.75% acrolein in 0.1 M phosphate buffer (PB; pH 7.4). Prostate tissues were removed and post-fixed at room temperature in 2% PFA and 2% acrolein in PB for 30 min. The prostate AP lobes were then placed in 2% osmium tetroxide in PB for 1 hour and subsequently washed in PB and dehydrated through a series of ethanol incubations, followed by propylene oxide, and incubated in 1:1 propylene oxide/Embed-812 (both from EMS) overnight. The next day, the prostate AP lobes were cut into distal and proximal parts and separately incubated in fresh Embed-812 for ∼2 hours. Next, the lobes were placed in the tips of plastic molds containing identifying labels, which were then filled with Embed-812. The molds were baked at 60°C for 3 days.

Ultrathin sections (∼70 nm) of the prostate AP lobes were cut using a Leica EM UCT6 ultratome (Leica Microsystems Inc.) and a Diatome diamond knife (EMS). The sections were collected on a 400-mesh thin-bar copper grids (T400-Cu, EMS) and counterstained with Uranyless^TM^ (EMS 22409) and lead citrate (EMS 22410). Images were captured at magnifications of 3,000× and 20,000× using a digital camera system (version 3.2, Advanced Microscopy Techniques, Woburn, MA, USA) attached to a Hitachi HT7800 transmission electron microscope (Hitachi High-Tech America, Inc.).

### Computational methods

#### scRNA-seq analysis

The raw data from 10X sequencing were processed using the Cell Ranger (*84*) (v.7, Genome build mm10) and Seurat (*85*) (v5.1.0) pipelines in R (v4.4.1) for all hashtag samples in this work. Only cells detected by both RNA and HTO (hashtag oligos) were included for downstream analyses. HTO signals were normalized using the centered log-ratio (CLR) transformation and used for cell demultiplexing, assigning each cell to an individual sample; cells labeled with multiple HTO barcodes or exhibiting negative global signal levels were excluded. To ensure high-quality data, we filtered out cells based on the following criteria: 1) >15% mitochondrial gene content, 2) <200 detected genes per cell, and 3) potential doublets identified using DoubletFinder (*86*) (v2.0.4). Subsequently, for downstream integration and analysis, raw count data were normalized and variance stabilized using the SCTransform function. Batch effects were corrected using Harmony (*87*) (v1.2.0) of all hashtag samples. Principal components were computed on the scaled data using the RunPCA function, and the first 30 principal components were used as input for cell clustering. Cell clusters were identified using the FindNeighbors function followed by the Louvain algorithm implemented in the FindClusters function, with a resolution parameter of 0.3. Finally, the RunUMAP function was used to generate a low-dimensional visualization of the scRNA-seq data, aiding in the identification of unbiased cell populations. Cell types were annotated based on canonical marker gene expression as follows: Luminal (*Epcam*, *Cd24a*, *Krt8*), Basal (*Epcam*, *Trp63*, *Krt5*), Macrophage (*Cd14*, *Aif1*), T cells (*Cd3e*, *Cd8a*), and Stromal cells (*Col5a2*, *Wnt2*, *Acta2*, *Fgf10*), using established references (*16, 18*).

Single-cell datasets (GSE146811) (*18*) from different time points following mouse castration were reanalyzed using the original analysis pipeline. For single-cell signature score analysis across four different timepoints, luminal cells from day3 after castration (WT Cas d3) and three additional timepoints from the above public single-cell datasets were selected: T00-Intact (WT Intact), T02_Cast_Day7 (WT Cas d7), and T04_Cast_Day28 (WT Cas d28); to maintain consistency with the original literature, we adopted the same cell annotation labels. For *Ptn* and *Igf1* related t-SNE visualizations, we also included the original t-SNE coordinates. To compare *Ptn* and *Igf1* expression changes across different stromal cell types between the intact time point and 7 days post-castration, stromal cells from all time points were reanalyzed using the Seurat pipeline. Batch effects were corrected using the CCAIntegration method applied to SCT-transformed normalized counts for these stromal cells. Cell clusters were identified using the FindNeighbors function, followed by clustering with the Louvain algorithm implemented in the FindClusters function, with the resolution parameter set to 0.3. Subsequently, clusters with negative *Ptn* or *Igf1* expression (median expression equal to zero) were excluded. Differential expression analysis was performed using the FindMarkers function with parameters min.pct = 0.2 and logfc.threshold = 0.2. Adjusted p-values obtained from the results were extracted and displayed at the top of the violin plots. For the GSEA analysis of cell death signatures between the intact timepoint and 7 days post-castration, the ferroptosis gene set was curated from a previous study (*28*), while the apoptosis, pyroptosis, and necroptosis gene sets were downloaded from the KEGG database (*88*); gene lists for each signature are provided in Table S1. For GSEA analysis comparing CSPC and CRPC patients, scRNA-seq data from tumor cells (GSE264573) (*37*) were reanalyzed using the original analysis pipeline. The avg_log2FC values generated by the FindMarkers function were used as rank-based input for GSEA analysis and visualization, performed using clusterProfiler (v4.8.3) (*89*).

#### Luminal cell type signature analysis and probability inference

To calculate distal luminal (LumA/Lum1) and proximal luminal (LumP/Lum2) gene signatures and infer prediction probabilities, we applied a method similar to that previously described by Karthaus and colleagues (*18*). To calculate distal/proximal luminal gene signature scores, genes were divided into 20 bins based on their mean normalized expression across all cells. These bins were defined by the distribution of expression for all genes. Gene expression values were centered, scaled, and transformed into the [0,1] range using a logistic function. For a gene set signature of *k* genes, the raw signature score for each cell was calculated as the mean normalized expression of the genes in the set. Marker genes for distal luminal cells (*e.g., Gsdma, Tgm4, Krt8*, *Cd24a, Pbsn, Hoxb13, Ceacam1, Prom1, Nkx3-1*) and proximal luminal cells (*e.g., Cldn10, Lrrc26, Ppp1r1b, Krt4, Wfdc2, Krt7, Tacstd2, Clu, Ly6a, Krt8*, *Cd24a, Ceacam1*) were previously identified (*16*). The raw signature scores were compared against a null distribution generated from 2,000 randomly selected gene sets, each containing *k* genes matched to the same expression bins as the original signature genes. The final per-cell activity score was derived by centering the raw score against the mean of the null distribution scores. For dot-plot visualization, per-cell signature scores were Z-scored by centering the values and scaling by the standard deviation. To infer signature changes after castration, we trained a boosted gradient decision tree classifier on intact prostate luminal cell states using XGBoost (v2.0.3)(*90*) in Python (v3.10.11) and applied the classifier to luminal cells from other time points and experiments. We then applied this classifier to luminal cells from other time points and experiments, using the prediction probability (ranging from 0 to 1) for color coding in the dotplot.

#### RNA-seq analysis

For mouse RNA-seq analysis, all bulk RNA-seq sequencing reads were aligned using STAR (version 2.7.11a) (*91*) to mouse reference sequence mm10. Gene expression values and TPM were quantitated with RSEM (version 1.3.1) (*92*). DESeq2 (v1.40.2) (*93*) was used for DEG identification, with genes with FDR <0.05 and |log2FC| > 1.5 identified as DEGs. All ligand-related genes from CellPhoneDB v5 (*94*) were selected for visualization in the volcano plot. Gene Ontology (GO) analysis was performed using g:Profiler (database version: Feb 12 2024) (*95*). For human RNA-seq analysis, TPM values for all enzalutamide responder or non-responder patients (*38*) were downloaded and used for GSEA analysis via clusterProfiler (v4.8.3). For comparison of the expression of MUFA/PUFA-phospholipid biosynthesis genes in human prostate cancer before and after ADT treatment, we analyzed an RNA-seq dataset (*46*) of prostate biopsies from seven patients with locally advanced or metastatic prostate cancer before and approximately 22 weeks after ADT initiation.

#### ChIP-seq data analysis

For ChIP-seq data analysis, mouse AR and NKX3-1 ChIP-seq datasets (GSE163145 (*44*) and GSE35971 (*43*), respectively) were used to investigate potential co-localization and binding in upstream genomic regions for *Gpx4*, *Mboat2*, *Acsl3*, *Scd1*, and *Fasn*. We also analyzed a human prostate tumor and adjacent normal tissue AR ChIP-seq dataset (GSE56288) (*45*), and a NKX3-1 ChIP-seq analysis of the LNCaP cell line (GSE40269) (*43*). Reads were aligned to the mouse reference genome (mm10) or human reference genome (hg38) using BWA-mem2 (v2.2.1) (*96*), and duplicate reads were marked using Picard (v3.3.0) (https://broadinstitute.github.io/picard/). To ensure data quality, reads were filtered using SAMtools (v1.17) (*97*), BEDTools (v2.31.0) (*98*), and BAMTools (v2.5.2) (*99*) based on the following criteria: 1) reads mapping to blacklisted regions; 2) reads that were marked as duplicates; 3) reads that were not marked as primary alignments; 4) reads that mapped to multiple locations; 5) reads containing > 4 mismatches. Filtered BAM files from each replicate were merged, and peaks were called using MACS2 (v2.2.9.1) (*100*) with default settings. ChIP-seq signal normalization and compression into reads-per-genomic-content (RPGC) BigWig files were performed using deepTools (v3.5.4) (*101*). Heatmap visualizations were generated using the computeMatrix function with the reference-point mode (--referencePoint center -b 3000 -a 3000) followed by the plotHeatmap function in deepTools. Genomic trait visualizations were created using GViz (1.50.0) (*102*).

### Statistical analysis

Statistical analyses were done using Prism (v.10, GraphPad Software). Statistical significance was determined using two-sided unpaired Student’s *t*-tests or Wilcoxon tests with Benjamini–Hochberg correction or FDR correction.

## Acknowledgements

We are grateful to Cory Abate-Shen, Christine Chio, Wei Gu, and Max Loda for critically reading the manuscript. We thank Caisheng Lu and Ching-Yuan Chen from the Flow Cytometry Core of the Columbia Center for Translational Immunology (CCTI) and Herbert Irving Comprehensive Cancer Center (HICCC), supported in part by awards S10OD020056, S10OD030282, and P30CA013696. We thank Erin Bush and Peter Sims in the Columbia Genomics and High Throughput Screening Shared Resource of the Herbert Irving Comprehensive Cancer Center, supported by the NIH/NCI Cancer Center Support Grant P30CA013696 and the National Center for Advancing Translational Sciences Support Grant UL1TR001873. We thank the Neuroanatomy Electron Microscopy Core at Weill Cornell Medicine for use of the Hitachi HT7800 transmission electron microscope obtained by the NIH Office of Research Infrastructure Program S10 Instrumentation Grant 1S10OD026974, and Ma’ayan Hirschkorn for technical assistance with EM procedures. We also thank Marie C. Hasselluhn and Tanner Dalton from Dr. Ken Olive’s lab (Columbia University Irving Medical Center) for assistance in organotypic cultures.

## Author contributions

W.L. and M.M.S. conceived and designed the overall study. W.L. designed and conducted most of the mouse experiments. W.L., J.B.Z. and Z.W dissociated the mouse prostate and J.B.Z. carried out all Western blot experiments. J.R.C. initiated the computational analysis, while Z.W. implemented most of it. S.X. assisted with mouse breeding and castration surgery. W.L., F.Z., and B.R.S. performed lipidomic analysis. W.L. and T.A.M. conducted the electron microscopy experiments. W.L., C.J.L., and H.H. carried out the human prostate cancer sample analysis. A.S.G. and J.P.C. generated conditional mouse models. M.M.S. supervised the experimental analyses. W.L. and M.M.S. wrote the initial manuscript with input from J.B.Z. and Z.W., and all authors contributed to its revision.

## Data and materials availability

The mouse bulk RNA-seq and scRNA-seq data have been deposited in the Gene Expression Omnibus database under accession numbers GSE295132 and GSE295388, respectively. Processed data and script are provided at Figshare link: 10.6084/m9.figshare.27984227.

## List of Supplementary Materials

Materials and Methods

Figures. S1 to S10

Tables S1 to S5

References (*74*-*102*)

---

**Fig. S1.**
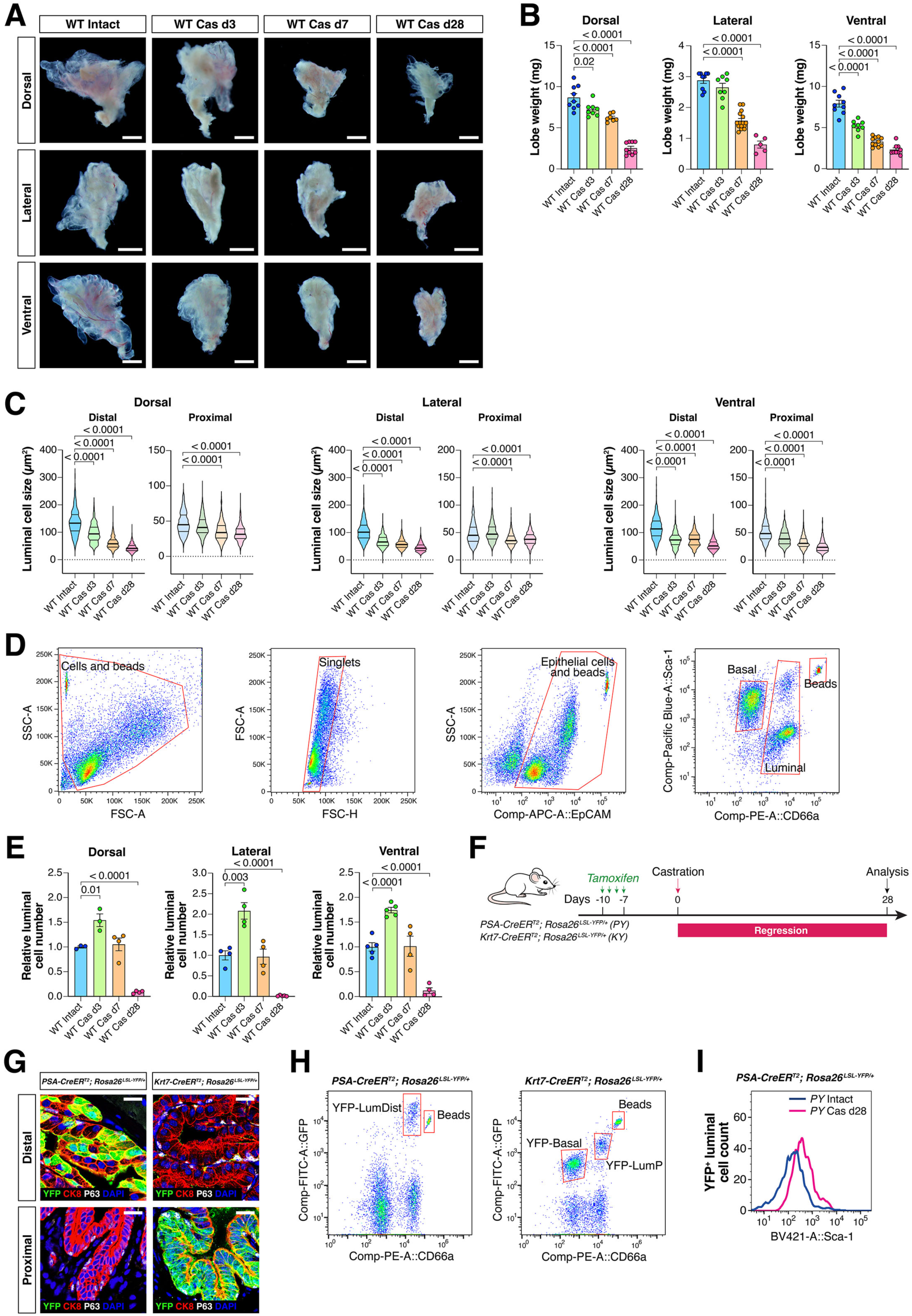
Features of prostate regression in wild type mice after castration. (**A**) Dark-field images of whole-mount dorsal (DP), ventral (VP), and lateral (LP) prostate lobes from 10-12 week-old wild-type (WT) mice that were hormonally-intact or castrated for the indicated number of days. Scale bars: 1 mm. (**B, C, E**) Prostate lobe weight (B), luminal cell size (C), and luminal cell number (E) in paired DP, VP, and LP lobes from intact and castrated WT mice. Distal luminal cells from AP, DP, VP, and LP lobes are named LumA, LumD, LumV, and LumL, respectively. (**D**) Flow sorting strategy for quantifying luminal cell number. (**F**) Schematic of lineage-tracing strategy for distal and proximal luminal cells after castration using tamoxifen-treated *PSA-CreER^T2^; Rosa26^LSL-YFP/+^* mice (*PY*) and *Krt7-CreER^T2^; Rosa26^LSL-YFP/+^* mice (*KY*), respectively. (**G**) Immunofluorescence staining of YFP, cytokeratin 8 (CK8), and p63 in AP lobes from *PY* and *KY* mice at day 10 after tamoxifen induction. Scale bars, 20 μm. (**H**) Flow sorting strategy for quantifying YFP-positive distal and proximal luminal cell numbers in paired AP lobes from *PY* and *KY* mice, respectively. (**I**) Staining intensity for Sca-1 in YFP-positive distal cells from *PY* mice that were hormonally-intact or 28 days after castration, respectively. All p-values were calculated using t-tests.

**Fig. S2.**
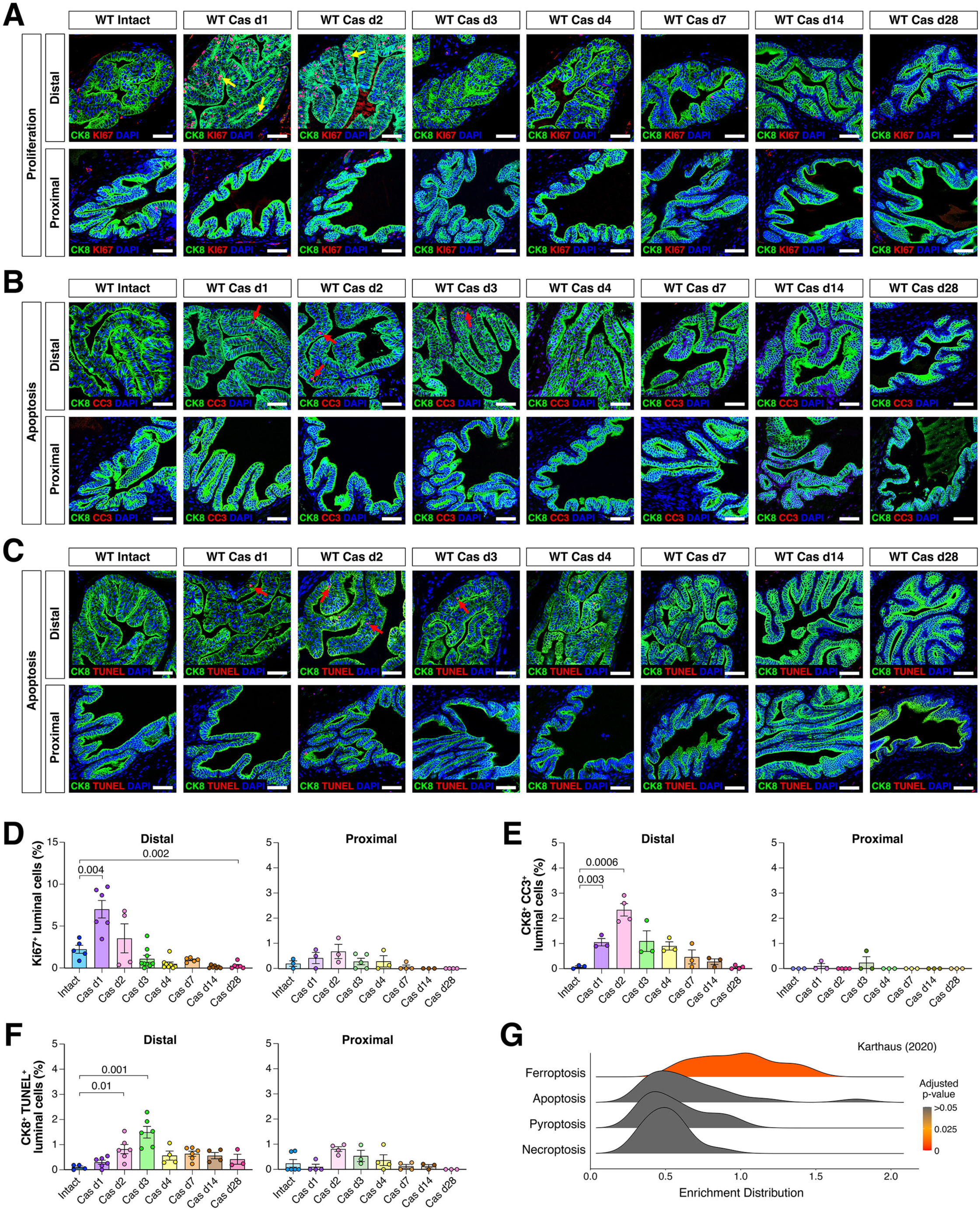
Time course of luminal cell proliferation and apoptosis in wild type mouse prostate after castration. (**A-C**) Immunofluorescence staining of Ki67, cytokeratin 8 (CK8), Cleaved Caspase-3 (CC3), and TUNEL to evaluate proliferation (A) or apoptosis (B-C) in distal and proximal luminal cells from AP lobes of WT mice that were either hormonally-intact or castrated for the indicated number of days. Scale bars, 50 μm. (**D-F**) Quantification of cell proliferation (D) or apoptosis (E-F) in distal and proximal luminal cells from WT mice; p-values were calculated by t-tests. (**G**) Ridgeline plots for the distribution of gene set enrichment (GSEA) results for four cell death signatures in luminal cells, comparing scRNA-seq data for WT Cas d7 versus WT intact prostate. Colors represent the adjusted p-values calculated by the clusterProfiler package.

**Fig. S3.**
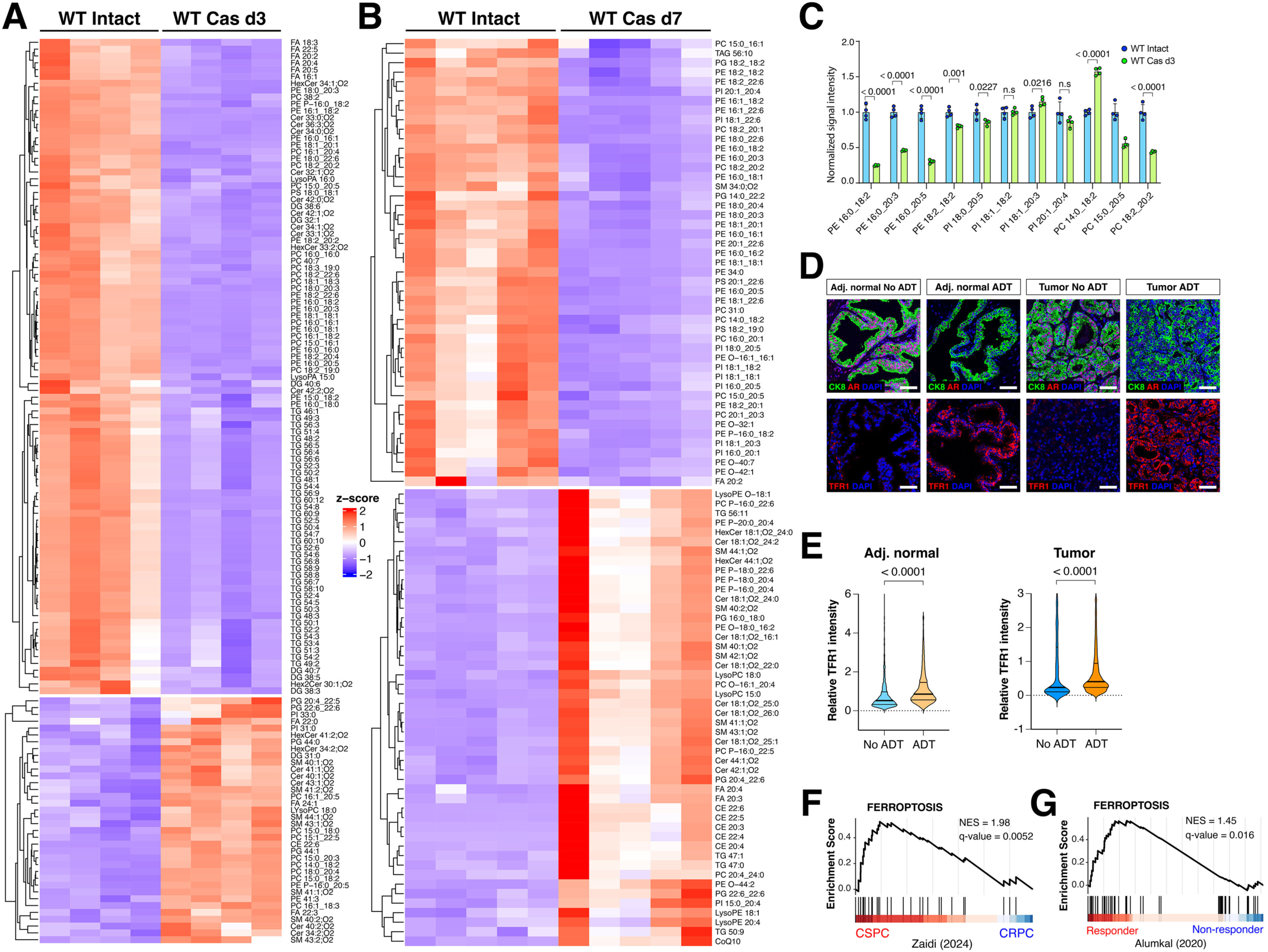
Castration induces ferroptosis in prostate luminal cells. (**A-B**) Heatmap of significantly changed lipid species (fold-change threshold = 1.5; FDR-corrected p < 0.05; each sample corresponds to lipids extracted from combined luminal cells from 5 mice) in WT intact mice versus WT Cas d3 mice (A), or WT intact mice versus WT Cas d7 mice (B). Each row represents z-score-normalized intensities of the detected lipid species. The relative abundance of each lipid is color-coded, with red indicating high signal intensity and blue indicating low signal intensity. CE, cholesteryl ester; Cer, ceramide; CoQ10, coenzyme Q10; FA, free fatty acid; SM, sphingomyelin; LysoPC, lysophosphatidylcholine; LysoPE, lysophosphatidylethanolamine; PE, Phosphatidylethanolamine; PC, phosphatidylcholine; PC O, ether-linked PC; PC P, plasmalogen PC; PE O, ether-linked PE; PE P, plasmalogen PE; PG, phosphatidylglycerol; PI, phosphatidylinositol; PS, phosphatidylserine; TAG, triacylglycerol; DG, Diacylglycerols; Hex2Cer, Dihexosylceramides. Lipids are annotated based on their fatty acyl compositions (for example, LysoPC 20:4 has 20 carbons and 4 double bonds) or as the sum of their total number of carbons and double bonds (for example, TAG 56:10 has a total of 56 carbons and 10 double bonds). (**C**) Abundance of representative poly-unsaturated fatty acid (PUFA)-phospholipids that are significantly downregulated in luminal cells from hormonally-intact and castrated WT mice at Cas d3, with each point corresponding to lipids extracted from combined luminal cells from 5 mice. (**D-E**) Immunofluorescence staining of CK8, AR, and TFR1 in benign prostate and prostate tumors from treatment-naive patients or patients who received neoadjuvant androgen-deprivation therapy (ADT) prior to transurethral resection or prostatectomy (D), with quantification of mean TFR1 staining intensity in CK8-positive adjacent normal and tumor cells (E). Scale bars, 20 µm. p-values were calculated using t-tests. (**F**) Gene Set Enrichment Analysis (GSEA) of scRNA-seq data(*37*) from CSPC and CRPC tumors following ADT. (**G)** GSEA using bulk RNA-seq(*38*) of biopsy samples from mCRPC patients who were responders or non-responders to enzalutamide.

**Fig. S4.**
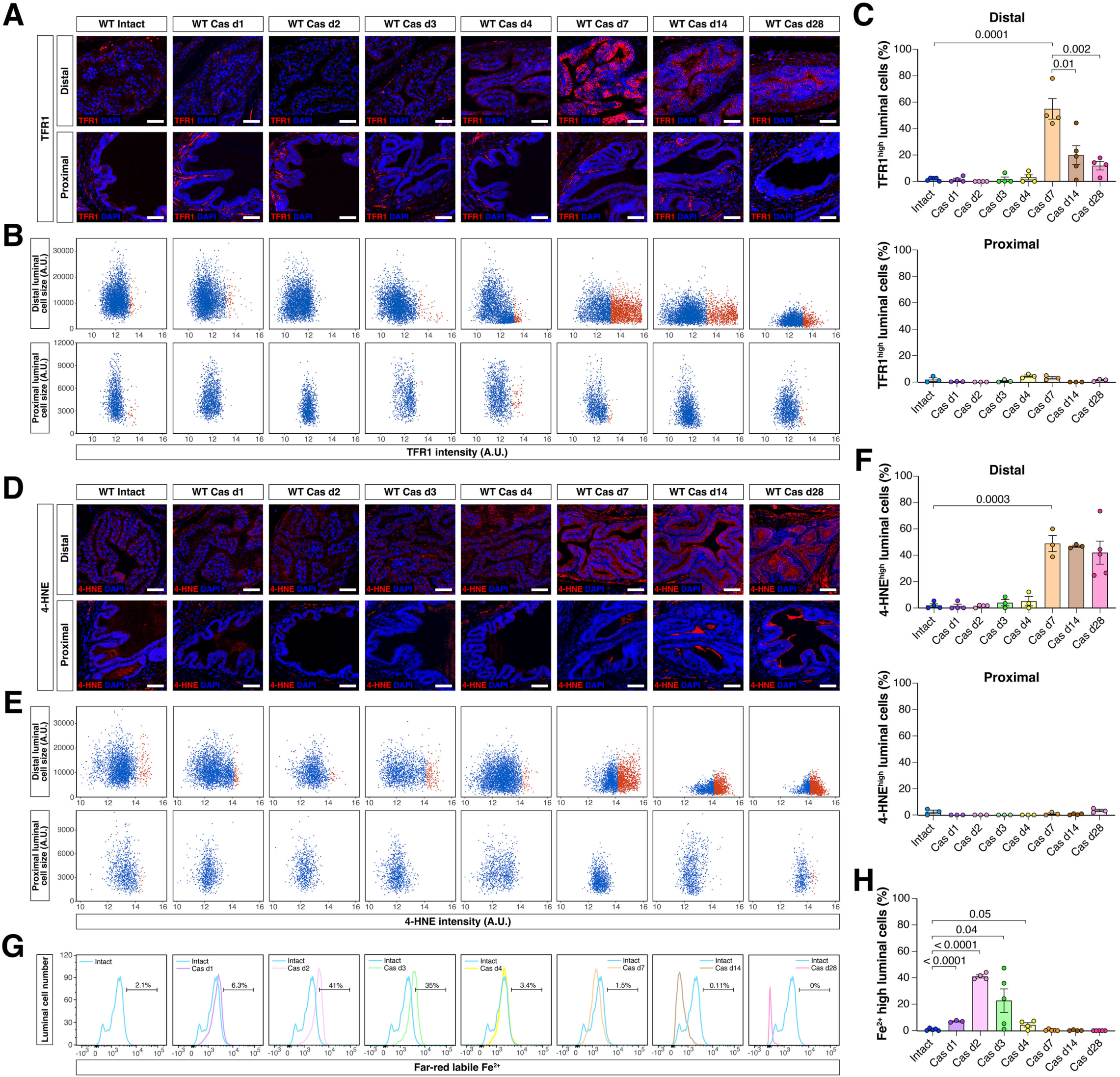
Time course of ferroptosis induction in wild type mouse prostate after castration. (**A** and **D**) Immunofluorescence staining of TFR1 (transferrin receptor) (A) or 4-HNE (4-hydroxynonenal) (D) in distal and proximal luminal cells from AP lobes of WT mice that were either hormonally-intact or castrated for the indicated number of days. Scale bars, 50 µm. (**B-C**) Quantitative image-based cytometry single-cell analysis (QIBC) of immunostained luminal cells in WT mice. Mean luminal TFR1 fluorescence intensity and CK8-positive luminal cell size from panel A are plotted in scatter diagrams (B). TFR1^high^ luminal cells are shown in red, and their ratio is quantified (C). (**E-F**) QIBC of immunostained luminal cells in WT mice. Mean luminal 4-HNE fluorescence intensity and CK8-positive luminal cell size from panel D are plotted in scatter diagrams (E). 4-HNE^high^ luminal cells are depicted in red, and their ratio is quantified (F). (**G-H**) Flow cytometry measurements of labile iron in luminal cells from WT mice using BioTracker FarRed dye (G), and quantification of the ratio of high labile iron luminal cells (H). All p-values were calculated using t-tests.

**Fig. S5.**
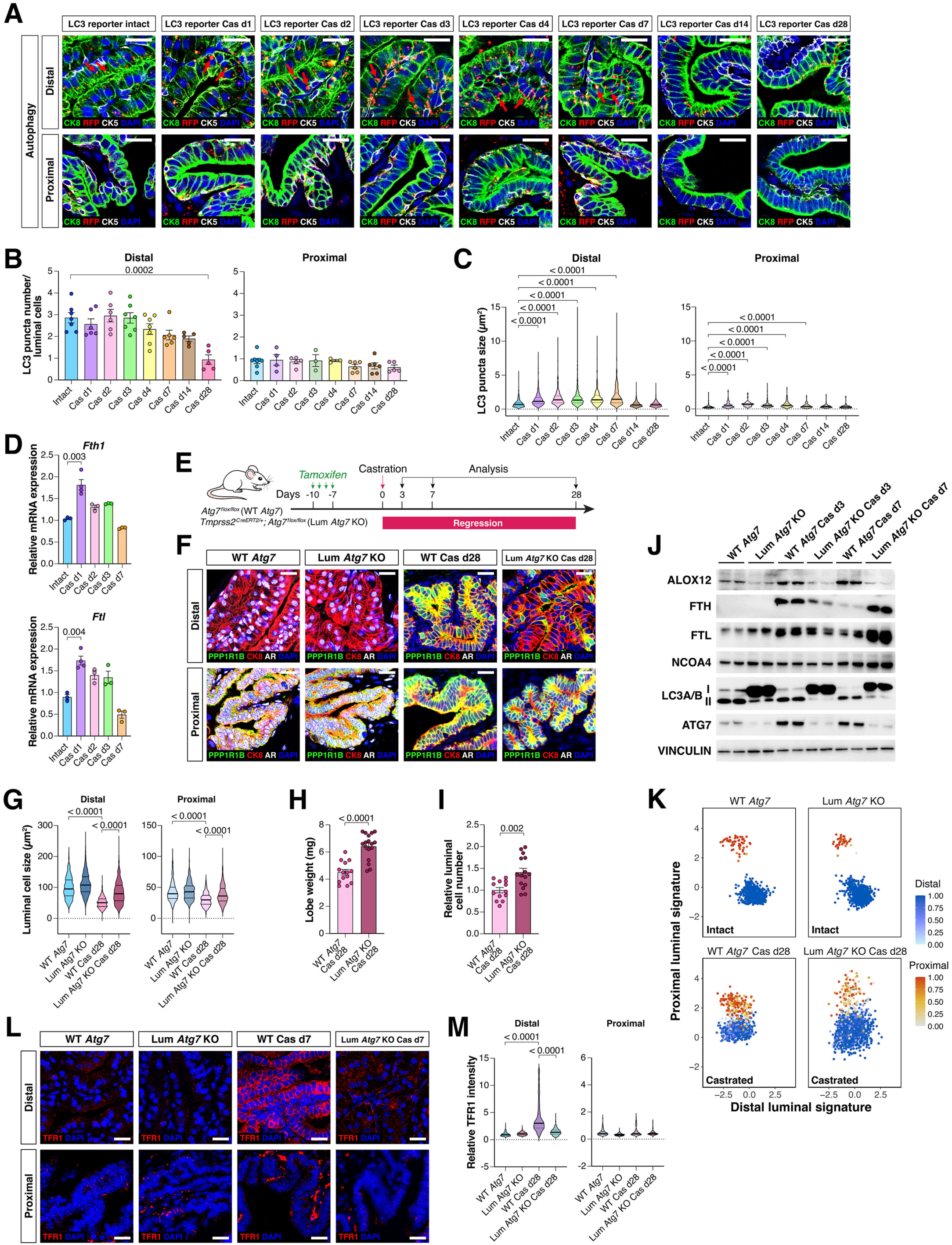
Castration-induced autophagy promotes ferroptosis induction in the prostate. (**A**) Immunofluorescence staining of RFP, cytokeratin 5 (CK5), and CK8 in AP lobes from LC3 reporter (*CAG-RFP-EGFP-LC3* transgenic) mice that were either hormonally-intact or castrated for the indicated number of days. (**B-C**) Quantification of LC3 puncta number (B) and puncta size (C) in distal and proximal luminal cells from LC3 reporter mice. (**D**) Quantitative PCR (qPCR) analysis of *Fth1* (encoding ferritin heavy chain) and *Ftl* (encoding ferritin light chain) expression. (**E**) Experimental strategy for inducible deletion of *Atg7* in luminal cells (Lum *Atg7* KO) in the absence or presence of castration. (**F**) Immunofluorescence staining of PPP1R1B, CK8, and AR and in AP lobes from WT *Atg7* and Lum *Atg7* KO mice that were hormonally-intact or castrated for 28 days. (**G-I**) Prostate luminal cell size (G), lobe weight (H), and luminal cell number (I) in AP lobes from mice of the indicated genotypes. (**J**) Western blotting for proteins associated with ferritinophagy and ferroptosis in luminal cells from WT *Atg7* and Lum *Atg7* KO mice, with each lane corresponding to combined protein extracted from 5 mice. (**K**) Scatterplots of scRNA-seq data for AP luminal cells from mice of the indicated genotypes, with each distal luminal (blue) or proximal luminal (red) cell assigned a distal (x-axis) and proximal luminal (y-axis) signature score. Color intensity corresponds to the probability strength of an XGBoost classifier assignment (color bar). (**L-M**) Immunofluorescence staining for TFR1 in mice of the indicated genotypes at day 7 after castration (**L**), and quantification of luminal TFR1 staining intensity in distal and proximal luminal cells (**M**). Scale bars in panel A, F, and L, 20 µm. All p-values were calculated using t-tests.

**Fig. S6.**
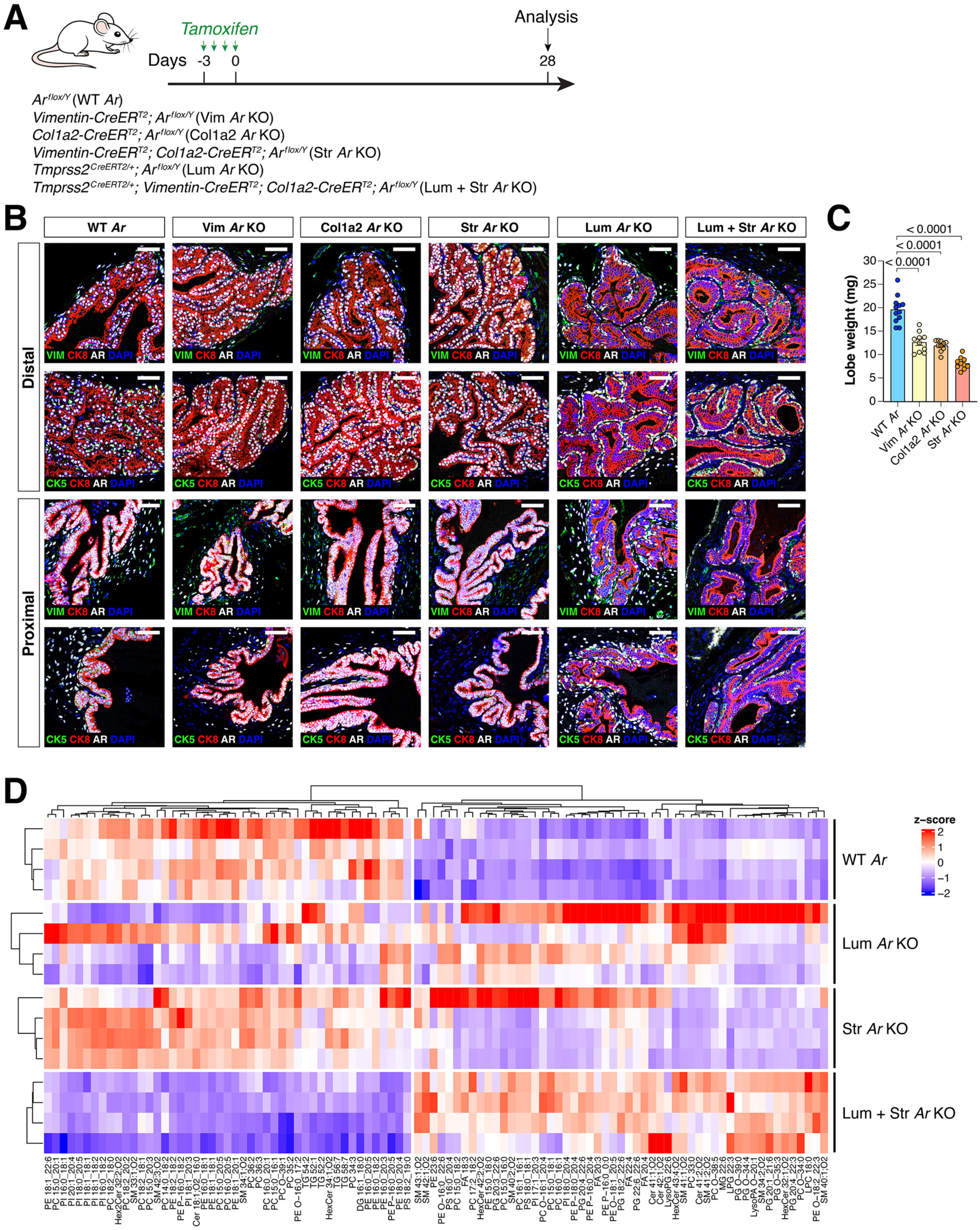
Intrinsic and extrinsic AR signaling regulate prostate luminal epithelial ferroptosis. (**A**) Experimental strategy and mouse genotypes for inducible deletion of AR in prostate luminal cells and/or stromal cells. (**B**) Immunostaining for AR, CK5, CK8, and Vimentin (VIM) in the AP lobes of the indicated genotypes to delete AR in luminal cells, stromal cells, or both at 28 days after tamoxifen-induction. Scale bars, 50 µm. (**C**) Prostate lobe weight in mice of the indicated genotypes at 28 days after tamoxifen treatment; p-values were calculated using t-tests. (**D**) Heatmap of significantly changed lipid species (fold-change threshold = 1.5; FDR-corrected P < 0.05; n = 4 biologically independent samples, with each sample corresponding to lipids extracted from combined luminal cells from 5 mice) in WT *Ar* mice versus Lum + Str *Ar* KO mice, along with the abundance of these lipid species in Lum *Ar* KO mice and Str *Ar* KO mice. Each column represents z-score-normalized intensities of the detected lipid species. The relative abundance of each lipid is color-coded, with red indicating high signal intensity and blue indicating low signal intensity.

**Fig. S7.**
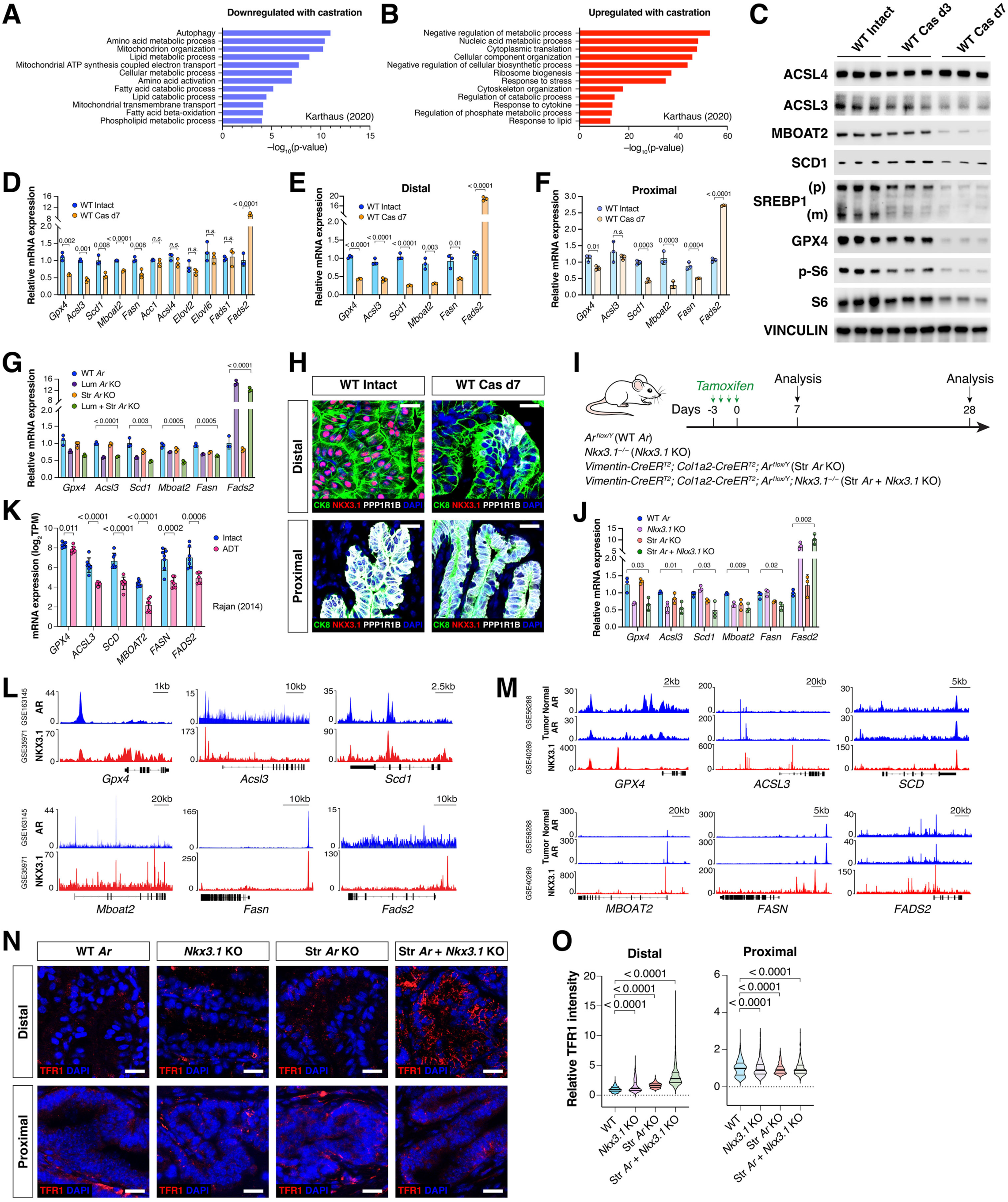
NKX3.1 mediates intrinsic AR signaling to suppress ferroptosis in distal luminal cells. (**A-B**) Gene Ontology (GO) analysis of enriched pathways for downregulated (A) and upregulated genes (B) in luminal cells from scRNA-seq of WT Cas d7 versus WT intact mouse prostate. (**C**) Western blotting of proteins involved in mTORC1 and MUFA/PUFA-phospholipid synthesis pathways in luminal cells from WT, Cas d3, and Cas d7 mice, with each lane corresponding to protein extracted from 5 mice combined. (**D-F**) qPCR analysis of the expression of *Gpx4* and genes regulating MUFA/PUFA-phospholipid synthesis in luminal cells (D), and in isolated distal luminal cells (E) or proximal luminal cells (F) from WT intact and Cas d7 mouse prostate. (**G**) qPCR analysis of the expression of *Gpx4* and genes regulating MUFA/PUFA-phospholipid synthesis in luminal cells from mice with induced deletion of *Ar* at day 7 after tamoxifen induction. (**H**) Immunostaining for NKX3.1, PPP1R1B, and CK8 in the AP lobes of WT intact and Cas d7 mice. Scale bars, 20 µm. (**I**) Experimental strategy and mouse genotypes for inducible AR deletion in prostate stromal cells in combination with *Nkx3.1* loss. (**J**) qPCR analysis of the expression of *Gpx4* and genes regulating MUFA/PUFA-phospholipid synthesis in luminal cells from mice with the indicated genotypes at 7 days after tamoxifen induction. (**K**) Plot of the expression of *GPX4* and genes regulating MUFA/PUFA-phospholipid synthesis in human prostate tumors, based on RNA-seq data (*46*) from treatment-naive patients or patients who have received neoadjuvant androgen-deprivation therapy (ADT). (**L-M**) AR and NKX3.1 ChIP-seq signal tracks near *Gpx4* and MUFA synthesis gene loci in mouse prostate datasets (*43, 44*) (L) and a human normal prostate and tumor dataset (*45*) (M). (**N**) Immunofluorescence staining of transferrin receptor (TFR1) in AP lobes of mice with the indicated genotypes at 7 days after tamoxifen induction. Scale bars, 20 µm. (**O**) Quantification of mean fluorescence staining intensity of TFR1 in distal and proximal luminal cells. All p-values were calculated using t-tests.

**Fig. S8.**
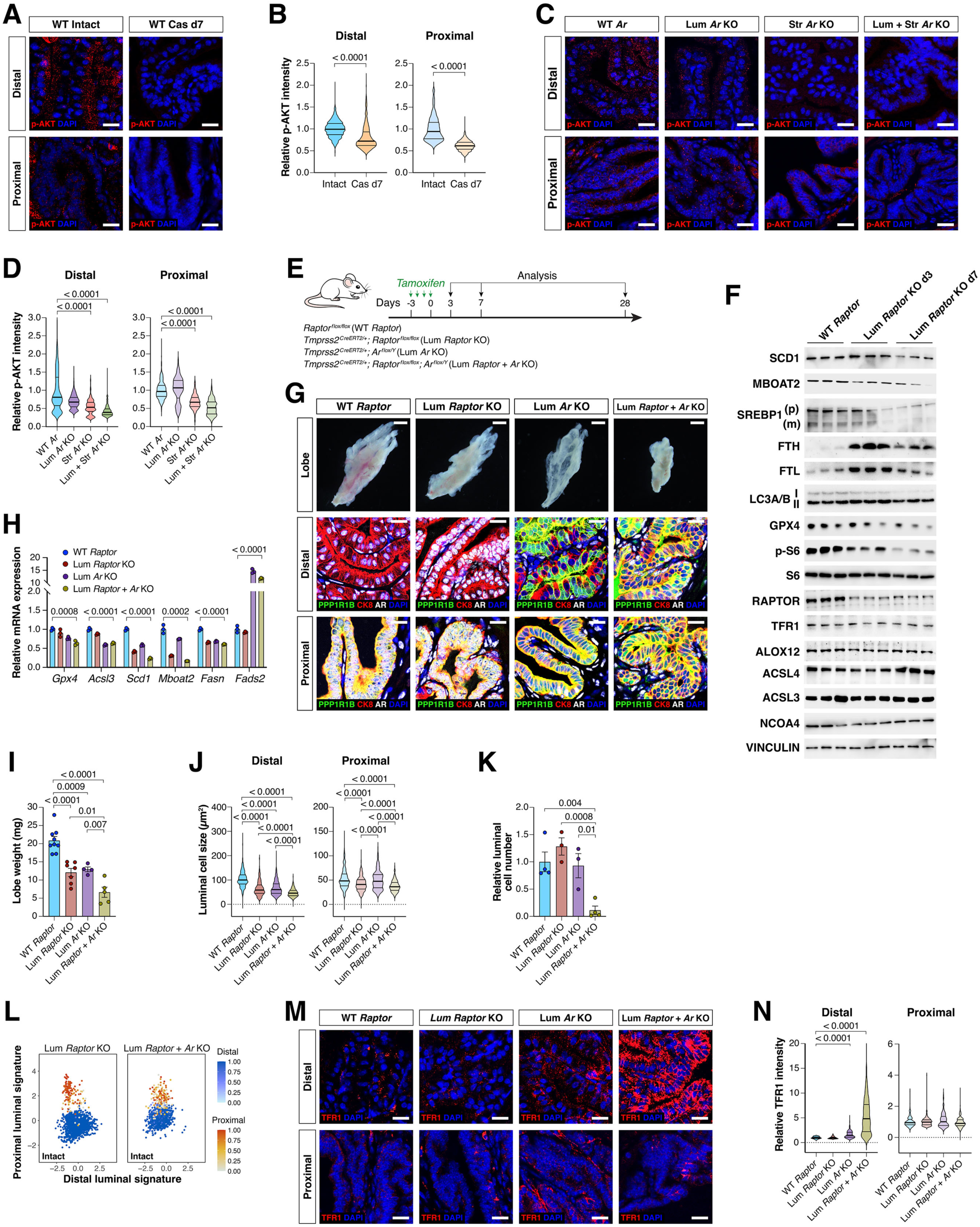
The mTORC1 pathway facilitates extrinsic AR signaling to suppress ferroptosis in distal luminal cells. (**A-B**) Immunofluorescence staining of p-AKT in AP sections from WT intact and WT Cas d7 mouse prostate (A), with quantification of mean p-AKT staining intensity in distal and proximal luminal cells (B). (**C-D**) Immunofluorescence staining of p-AKT in AP sections from mice of the indicated genotypes at 7 days after tamoxifen induction (C), with quantification of mean p-AKT staining intensity in distal and proximal luminal cells (D). (**E**) Experimental strategy and genotypes for inducible *Raptor* and/or *Ar* deletion in prostate luminal cells. (**F**) Western blotting of GPX4 and MUFA- and PUFA-phospholipid synthesis-associated proteins in luminal cells from inducible *Raptor* deletion at 3 and 7 days after tamoxifen induction. (**G**) Dark-field images of AP lobes and immunofluorescence staining of AP sections from control mice and mice with induced deletion of *Ar* in luminal cells (Lum *Ar* KO), *Raptor* in luminal cells (Lum *Raptor* KO), or both (Lum *Raptor* + *Ar* KO), at day 28 after tamoxifen induction. Scale bars for whole-mounts, 1 mm; for sections, 20 µm. (**H**) qPCR analysis of the expression of *Gpx4* and genes regulating MUFA/PUFA-phospholipid synthesis in luminal cells from mice of the indicated genotypes at 7 days after tamoxifen induction. (**I-K**) Lobe weight (I), luminal cell size (J), and luminal cell number (K) from mice of the indicated genotypes at day 28 after tamoxifen induction. (**L**) Scatterplots of scRNA-seq data showing transcriptomic changes in distal (blue) and proximal (red) luminal cells from control mice or mice with induced *Raptor or Raptor + Ar* deletion at day 28 after tamoxifen induction. (**M-N**) Immunofluorescence staining of TFR1 in AP lobes from mice with the indicated genotypes at 7 days after tamoxifen induction (M), with quantification of mean TFR1 staining intensity in distal and proximal luminal cells (N). All p-values were calculated using t-tests.

**Fig. S9.**
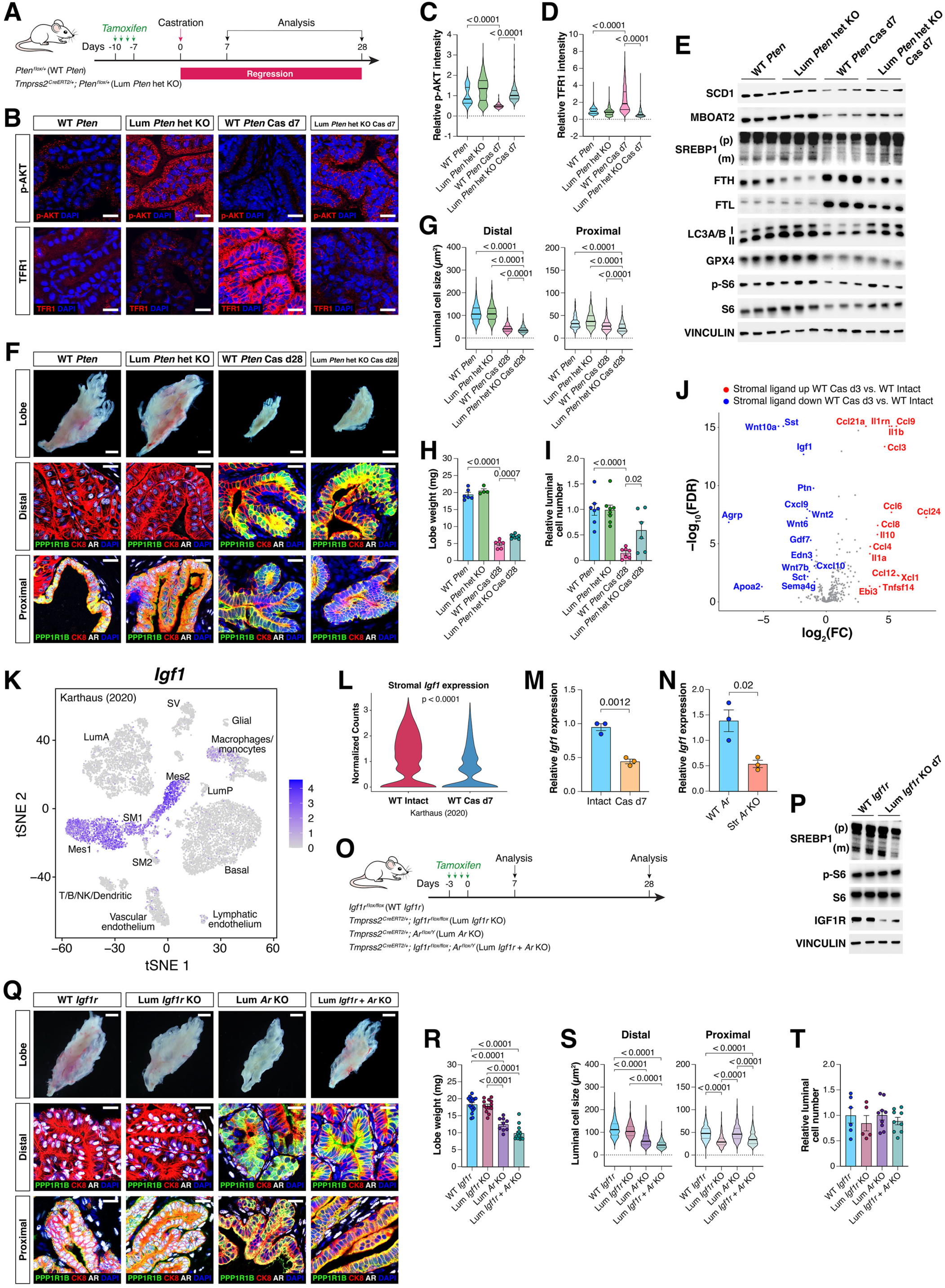
IGF1 does not mediate extrinsic AR signaling to suppress ferroptosis induction. (**A**) Strategy and genotypes for inducible deletion of a single *Pten* allele in prostate luminal cells. (**B-D**) Immunofluorescence staining of p-AKT and TFR1 in AP sections from mice of the indicated genotypes at 7 days after tamoxifen induction, with quantification of mean p-AKT staining intensity (C) and TFR1 (D) in distal luminal cells. (**E**) Western blotting of MUFA-phospholipid synthesis-associated proteins and mTORC1 pathway proteins in luminal cells from the indicated genotypes at 7 days after castration. (**F**) Dark-field whole-mount images of AP lobes and immunofluorescence staining of AP sections from mice with the indicated genotypes at 28 days after castration. Scale bars for whole-mounts, 1 mm, for sections, 20 µm. (**G-I**) Plots of AP luminal cell size (G), lobe weight (H), and luminal cell number (I) from mice of the indicated genotypes at 28 days after castration. (**J**) Volcano plot displaying significantly altered prostate stromal ligands from RNA-seq analyses of stromal cells from WT-intact and 3-day castrated mice. (**K**) t-SNE plot shows *Igf1* expression in scRNA-seq (*18*) of the WT intact mouse prostate. (**L**) Violin plot illustrates normalized *Igf1* expression levels in stromal cells from scRNA-seq of the WT intact and Cas d7 prostate. (**M-N**) qPCR analysis of the expression of *Igf1* in stromal cells from mice with the indicated genotypes. (**O**) Experimental strategy and mouse genotypes for inducible *Ar* deletion in prostate stromal cells in combination with *Igf1r* deletion. (**P**) Western blotting of IGF1R and mTORC1 pathway proteins in luminal cells from the indicated genotypes at 7 days after tamoxifen treatment. (**Q**) Dark-field whole-mount images of AP lobes and immunofluorescence staining of AP sections from mice with the indicated genotypes at 28 days after tamoxifen treatment. Scale bars for whole-mounts, 1 mm, for sections, 20 µm. (**R-T**) Plots of AP lobe weight (R), luminal cell size (S), and luminal cell number (T) from mice of the indicated genotypes at 28 days after castration. All p-values were calculated using t-tests.

**Fig. S10.**
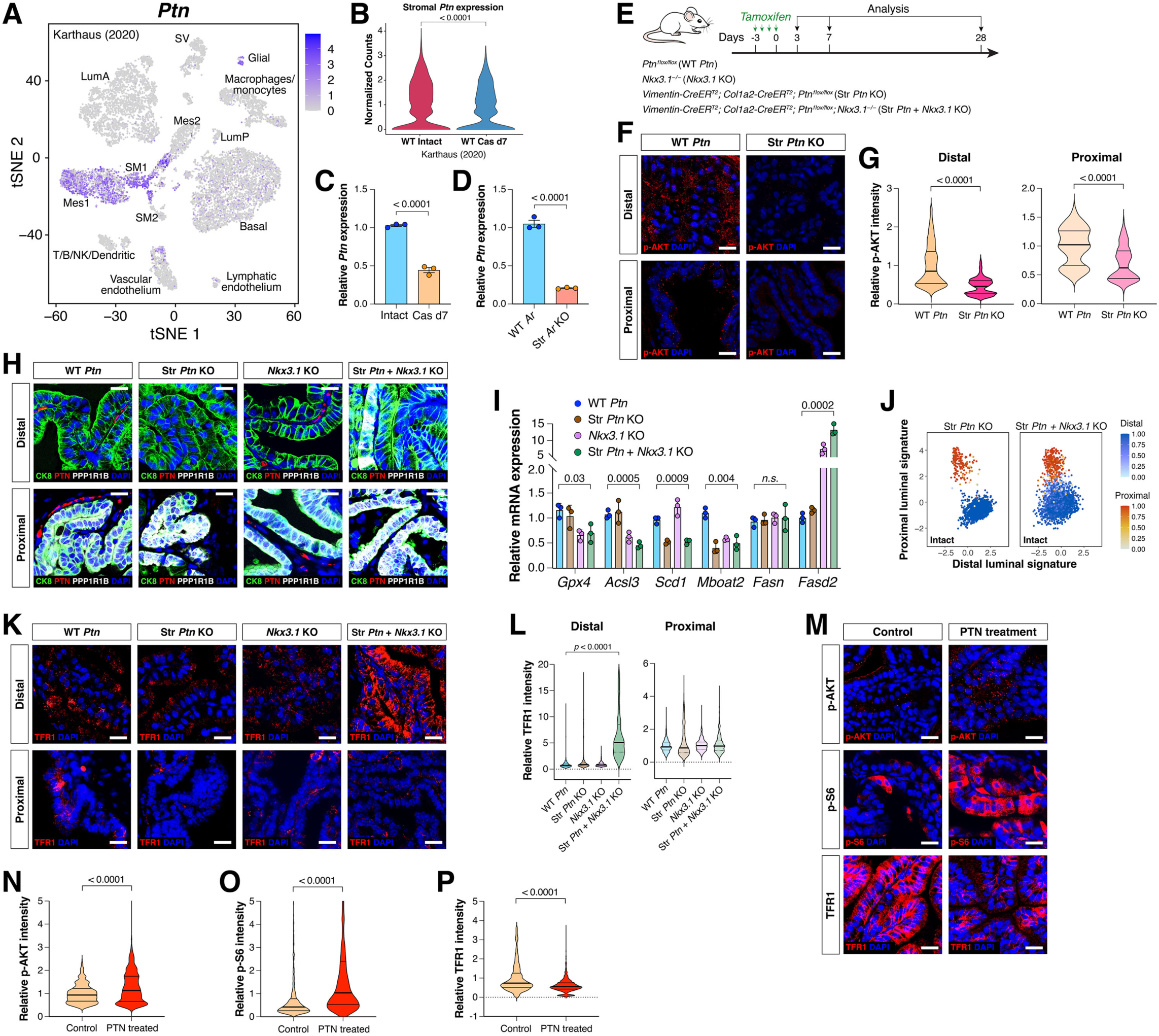
PTN mediates extrinsic AR signaling to suppress ferroptosis induction. (**A**) t-SNE plot shows *Ptn* expression in scRNA-seq (*18*) of the WT intact mouse prostate. (**B**) Violin plot illustrates normalized *Ptn* expression levels in stromal cells from scRNA-seq of the WT intact and Cas d7 prostate. (**C-D)** qPCR analysis of *Ptn* expression in stromal cells from mice with the indicated genotype and treatment. (**E**) Experimental strategy and mouse genotypes for inducible *Ptn* deletion in prostate stromal cells in combination with *Nkx3.1* loss. (**F-G**) Immunofluorescence staining of p-AKT in AP sections from WT control and Str *Ptn* KO mouse prostate at day 7 after tamoxifen treatment (F), with quantification of mean p-AKT staining intensity in distal and proximal luminal cells (G). (**H**) Immunofluorescence staining of CK8, PTN, and PPP1R1B in AP sections from mice with indicated genotypes at day 28 after tamoxifen treatment. (**I**) qPCR analysis of the expression of *Gpx4* and genes regulating MUFA/PUFA-phospholipid synthesis in luminal cells from mice with the indicated genotypes at 7 days after tamoxifen induction. (**J**) Scatterplots of scRNA-seq data showing transcriptomic changes in distal (blue) and proximal (red) luminal cells from control mice or mice with induced *Ptn or Ptn + Nkx3.1* deletion at day 28 after tamoxifen induction. (**K-L**) Immunofluorescence staining of TFR1 in AP lobes from mice with the indicated genotypes at 7 days after tamoxifen induction (K), with quantification of mean TFR1 staining intensity in distal and proximal luminal cells (L). (**M-P**) Immunofluorescence staining of p-AKT, p-S6, and TFR1 in AP lobes from WT mice treated with recombined mouse PTN protein or PBS control at day 7 after castration (M), with quantification of mean p-AKT staining intensity (N), mean p-S6 staining intensity (O), and TFR1 staining intensity in distal luminal cells (P). All p-values were calculated using t-tests.

**Table S1.**
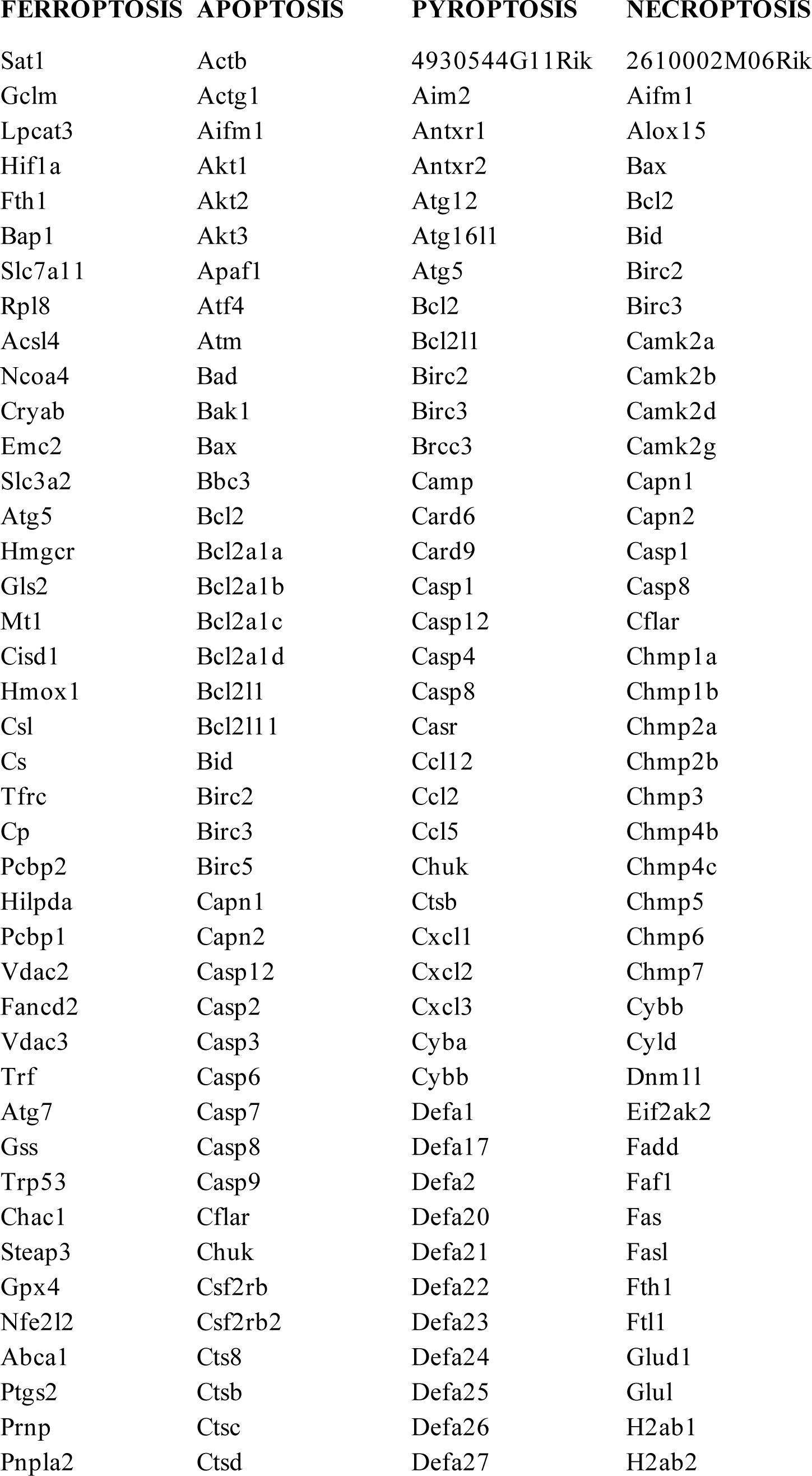
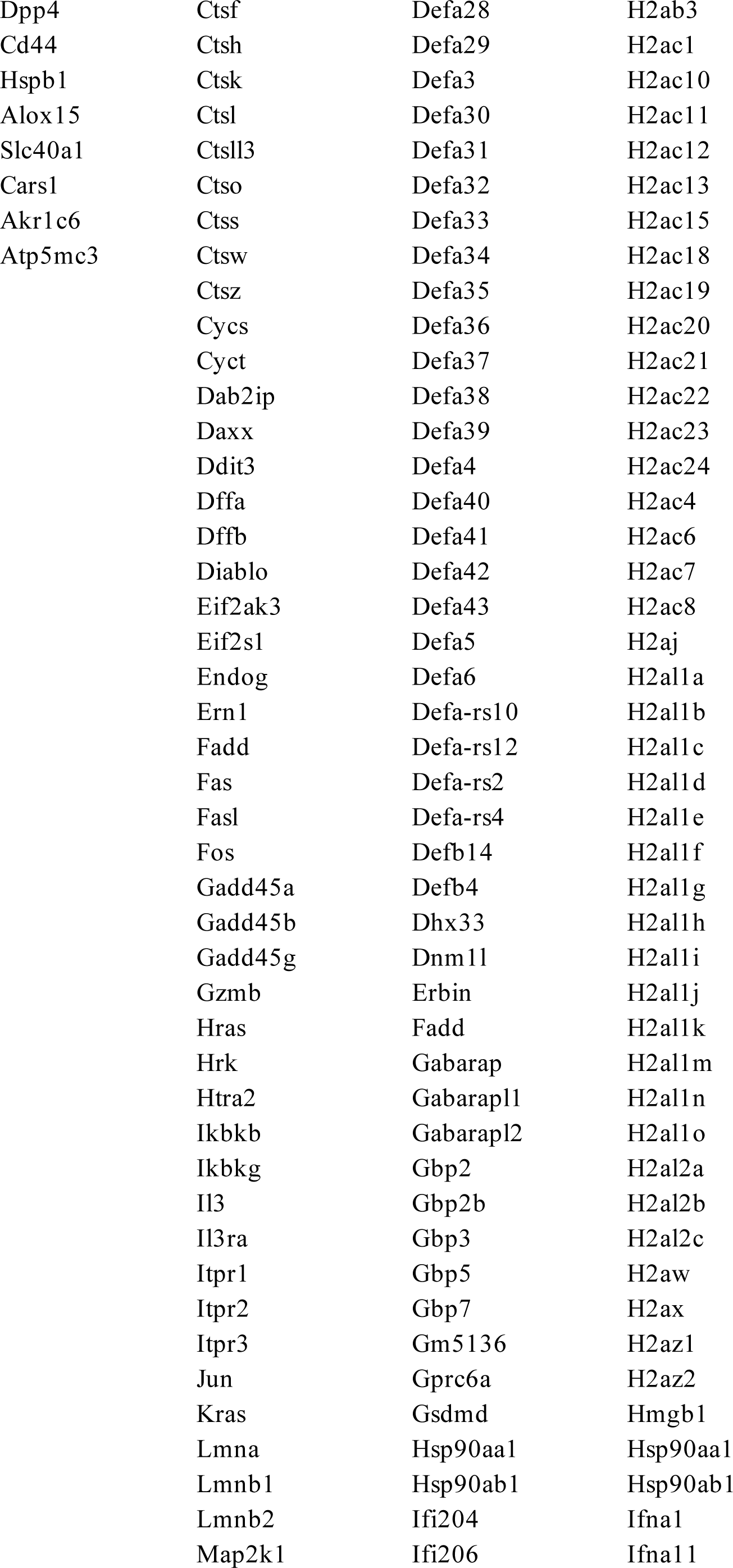
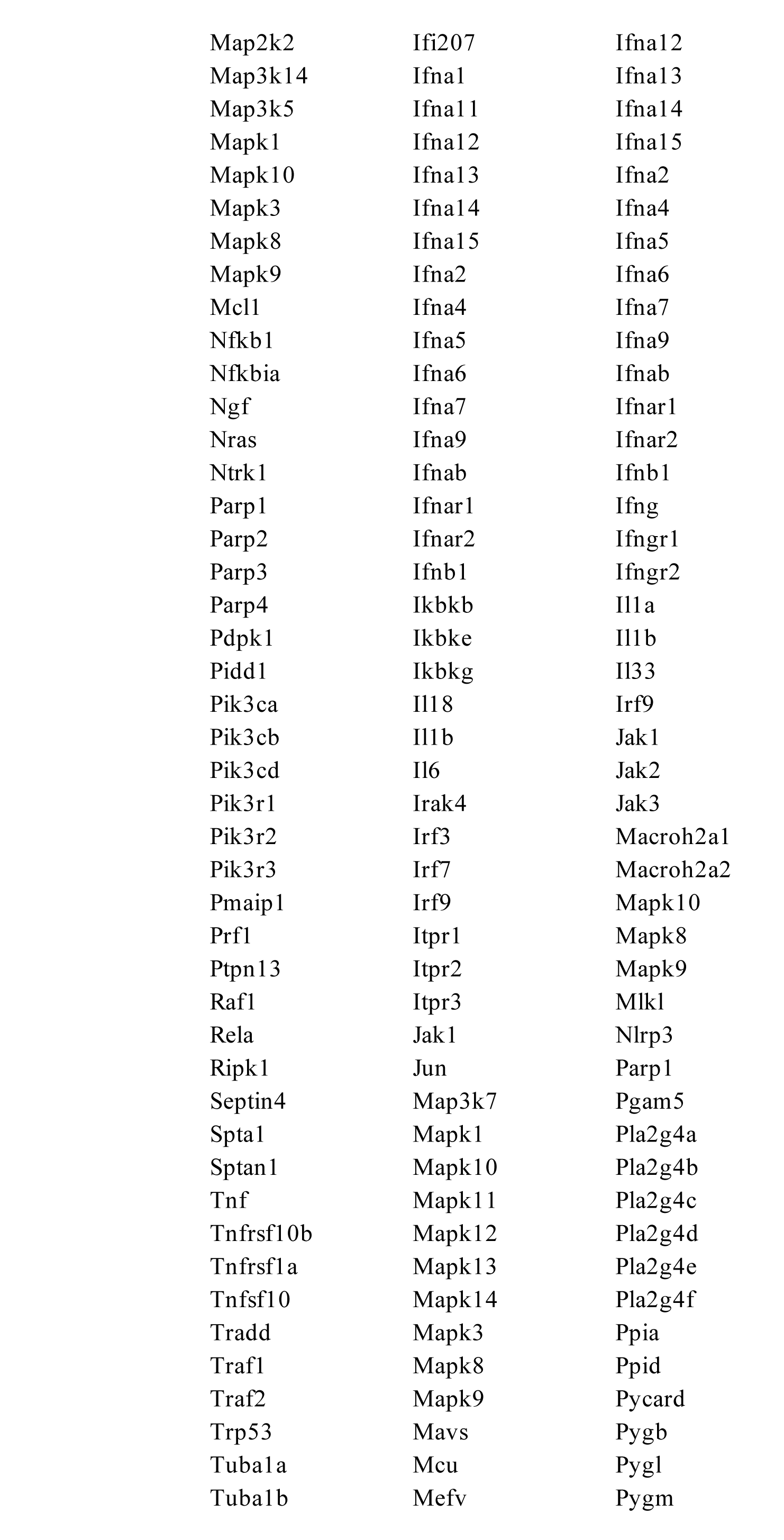
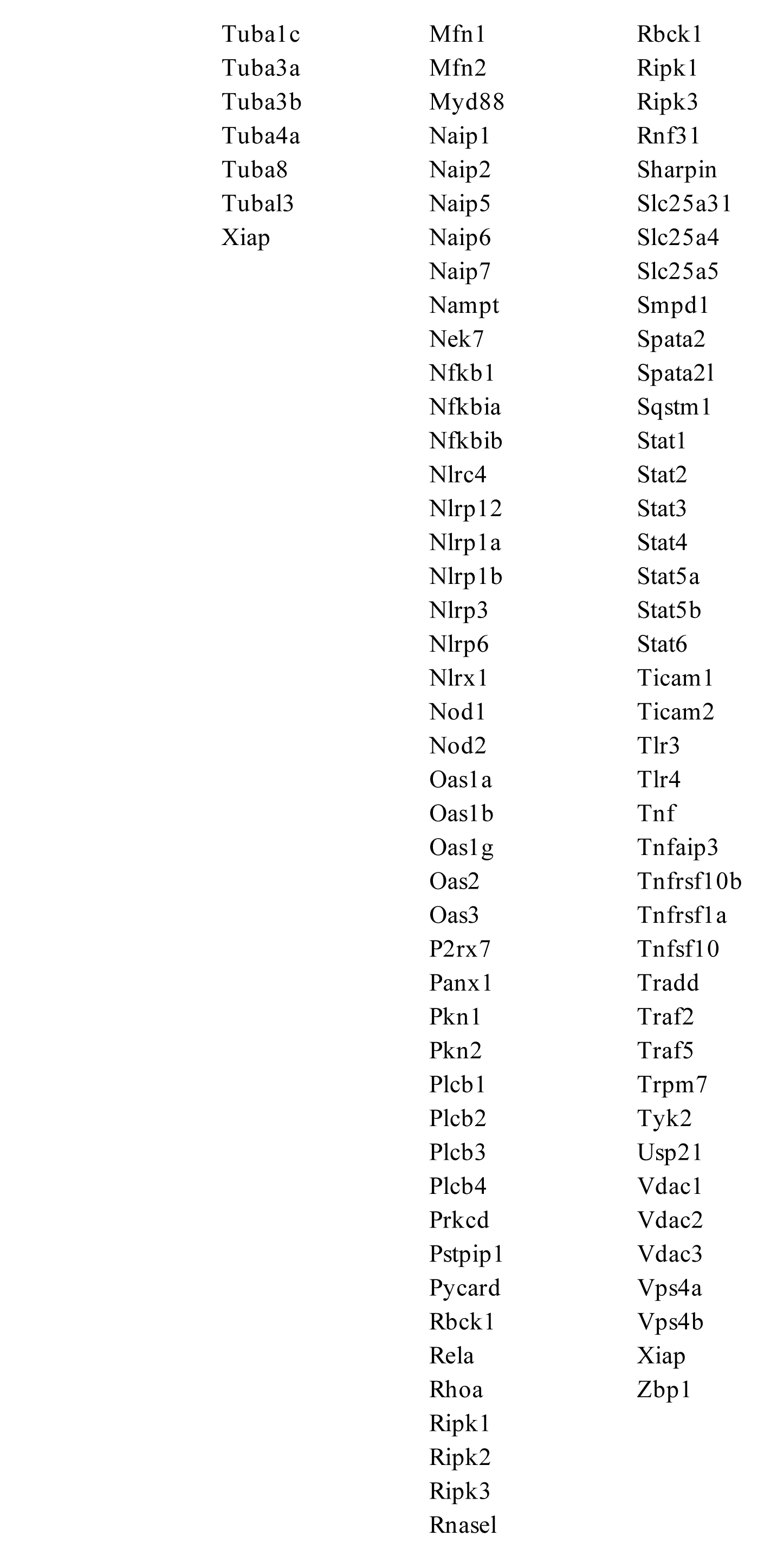
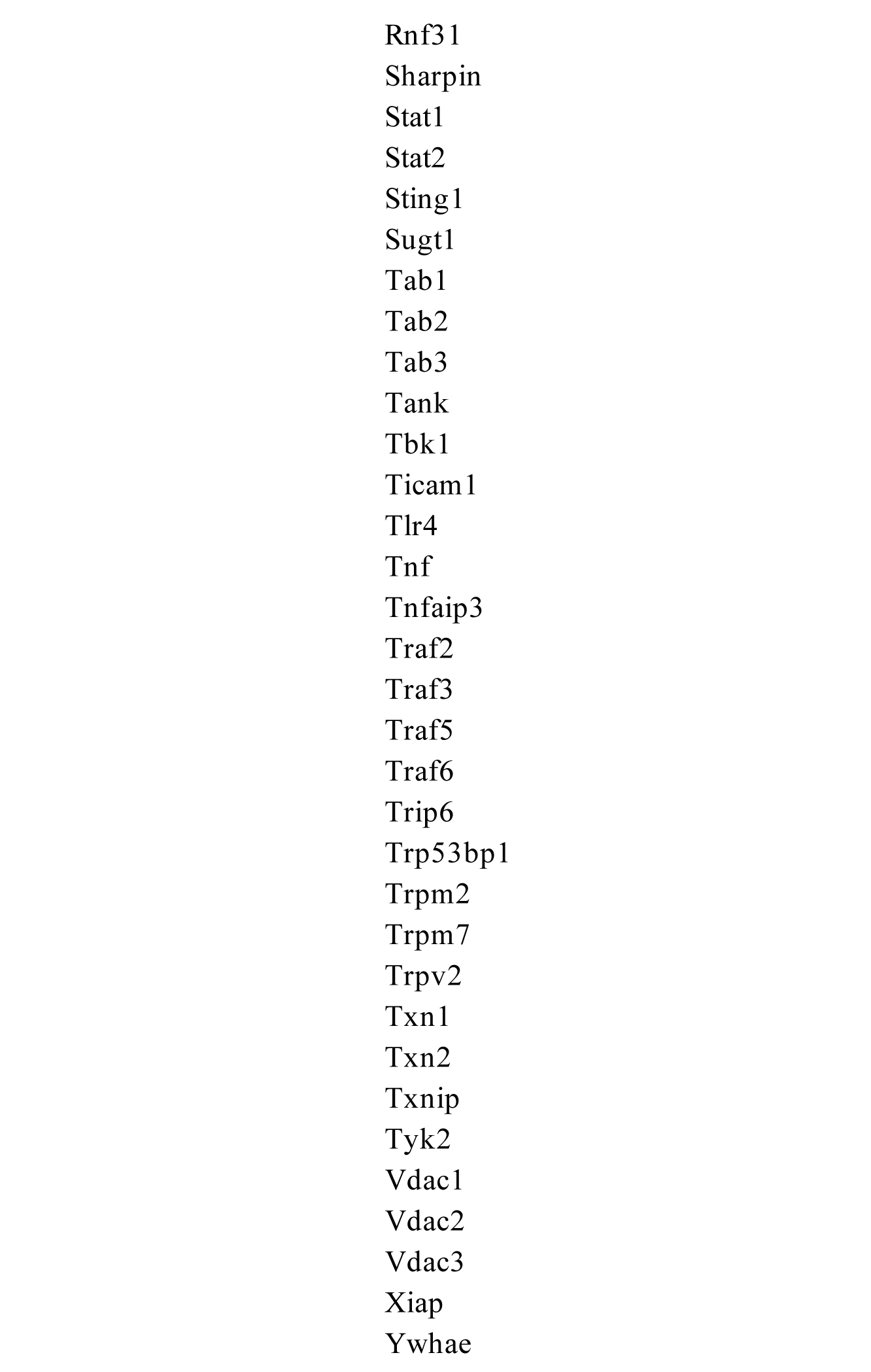
Gene lists for cell death signatures.

**Table S2.** Clinical parameters of human primary prostate cancer patients receiving neoadjuvant ADT and treatment-naïve controls.

| ID Code | Procedure type | Type of pre-procedure treatment | Age at Procedure | Race | Ethnicity | Pathological features of tumor |  |  |
| --- | --- | --- | --- | --- | --- | --- | --- | --- |
|  |  |  |  |  |  | Gleason score | T-stage | N-Stage |
| P1 | TURP | Docetaxel, Zoladex, Eligard, Enzalutamide | 80 | White | Non Hispanic | N/A | N/A | N/A |
| P2 | TURP | Abiraterone, Eligard | 52 | White | Non Hispanic | N/A | N/A | N/A |
| P3 | RALP | Degarelix, Apalutamide | 47 | White | Non Hispanic | GG5 | pT3b | pN0 |
| P4 | RALP | Lupron | 65 | White | Non Hispanic | GG3 | pT3a | pN0 |
| P5 | RALP | Degarelix | 67 | White | Non Hispanic | GG5 | pT3b | pN1 |
| P6 | RALP | Degarelix | 71 | White | Non Hispanic | GG5 | pT2 | pN0 |
| P7 | RALP | Degarelix | 71 | Other Combinations Not Described | Hispanic | GG5 | pT3b | pN1 |
| P8 | RALP | Degarelix | 65 | Other Combinations Not Described | Non Hispanic | GG3 | pT2 | pN0 |
| U1 | RALP | No treatment | 60 | White | Hispanic | GG5 | pT3b | pN1 |
| U2 | RALP | No treatment | 65 | White | Hispanic | GG5 | pT3b | pN1 |
| U3 | RALP | No treatment | 59 | Unknown | Unknown | GG4 | pT2 | pN0 |
| U4 | RALP | No treatment | 70 | White | Unknown | GG2 | pT2 | pN0 |
| U5 | RALP | No treatment | 51 | Other Combinations Not Described | Hispanic | GG5 | pT3b | pN1 |
| U6 | RALP | No treatment | 66 | Other Combinations Not Described | Hispanic | GG5 | pT3b | pN0 |
| U7 | RALP | No treatment | 75 | White | Non Hispanic | GG5 | pT3a | pN0 |
| U8 | RALP | No treatment | 66 | Other Combinations Not Described | Non Hispanic | GG3 | pT3b | pN1 |
| U9 | RALP | No treatment | 74 | White | Non Hispanic | GG4 | pT3a | pN0 |
| U10 | RALP | No treatment | 60 | Black | Non Hispanic | GG5 | pT3a | pN0 |
| U11 | RALP | No treatment | 60 | Other Combinations Not Described | Hispanic | GG5 | pT3a | pN0 |
| U12 | RALP | No treatment | 64 | White | Non Hispanic | GG3 | pT2 | pN0 |

RALP, Radical prostatectomy; TURP, transurethral resection of prostate; GG, Gleason grade

**Table S3.** Clinical parameters of human primary prostate tumor samples used for organotypic culture assay.

| ID Code | Procedure type | Type of pre-procedure treatment | Age at procedure | Race | Ethnicity | Pathological features of tumor t |  |  |
| --- | --- | --- | --- | --- | --- | --- | --- | --- |
|  |  |  |  |  |  | Gleason score | T-stage | N-Stage |
| #1 | RALP | No treatment | 69 | Black | Non Hispanic | GG2 | pT3b | pN0 |
| #5 | RALP | No treatment | 71 | White | Non Hispanic | GG5 | pT3b | pN1 |
| #7 | RALP | No treatment | 55 | Black | Non Hispanic | GG3 | pT3a | pN1 |

RALP, Radical prostatectomy; GG, Gleason grade

**Table S4.**
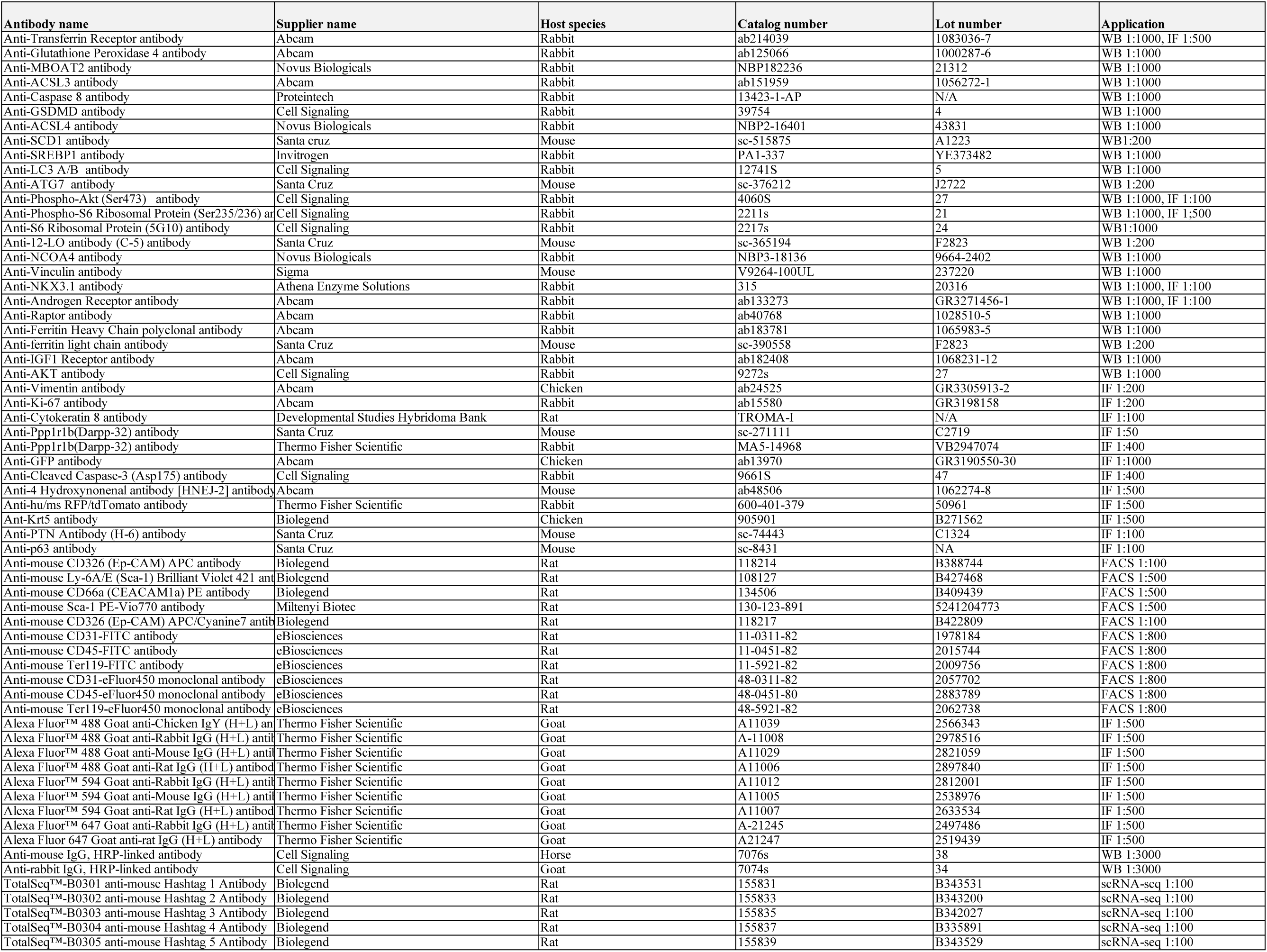
Antibodies used in this study.

**Table S5.**
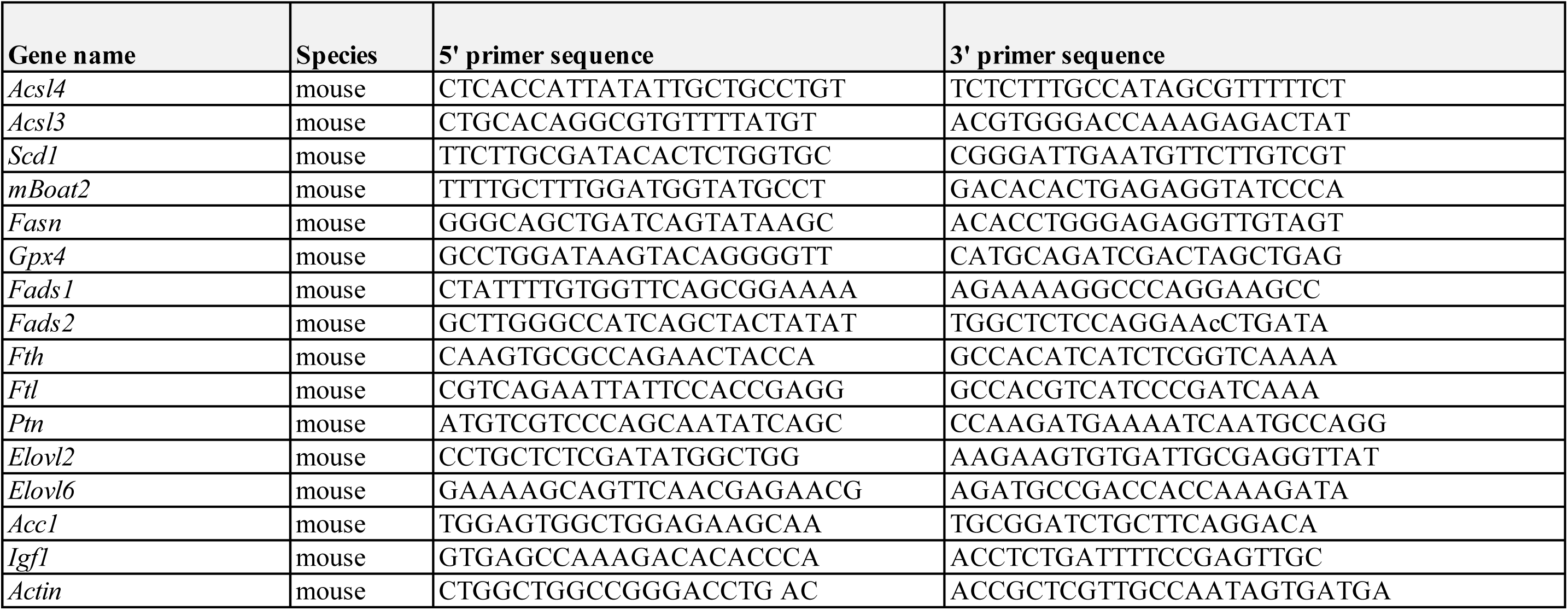
qPCR primers used in this study.

